# Pan-cancer in silico analysis of somatic mutations in G-protein coupled receptors: The effect of evolutionary conservation and natural variance

**DOI:** 10.1101/2021.10.25.465693

**Authors:** B.J. Bongers, M. Gorostiola González, X. Wang, H.W.T. van Vlijmen, W. Jespers, H. Gutiérrez-de-Terán, K. Ye, A.P. IJzerman, L.H. Heitman, G.J.P. van Westen

**Affiliations:** Division of Drug Discovery and Safety, Leiden Academic Centre for Drug Research, Leiden University, Leiden, The Netherlands; ONCODE Institute, Leiden, The Netherlands; Janssen Pharmaceutica NV, Beerse, Belgium; Department of Cell and Molecular Biology, Uppsala University, Uppsala, Sweden; School of Electronic and Information Engineering, Xi’an Jiaotong University, Xi’an, China

**Keywords:** Pareto Optimization, mutations, multi-objective, GPCR, cancer, GDC, natural variance, 1000 Genomes

## Abstract

G protein-coupled receptors (GPCRs) form the most frequently exploited drug target family, moreover they are often found mutated in cancer. Here we used an aggregated dataset of mutations found in cancer patient samples derived from the Genomic Data Commons and compared it to the natural human variance as exemplified by data from the 1000 Genomes project. While the location of these mutations across the protein domains did not differ significantly in the two datasets, a mutation enrichment was observed in cancer patients among conserved residues in GPCRs such as the “DRY” motif. We subsequently created a ranking of high scoring GPCRs, using a multi-objective approach (Pareto Front Ranking). The validity of our approach was confirmed by re-discovery of established cancer targets such as the LPA and mGlu receptor families, and we identified novel GPCRs that had not been directly linked to cancer before such as the P2Y Receptor 10 (*P2RY10*). As a proof of concept, we projected the structurally investigated mutations in the crystal structure of the C-C Chemokine (CCR) 5 receptor, one of the high-ranking GPCRs previously linked to cancer. Several positions were pinpointed that relate to either structural integrity or endogenous and synthetic ligand binding, providing a rationale to their mechanism of influence in cancer. In conclusion, this study identifies a list of GPCRs that are prioritized for experimental follow up characterization to elucidate their role in cancer. The computational approach here described can be adapted to investigate the roles in cancer of any protein family.

**Author summary:** Despite cancer being one of the most studied diseases due to its high mortality rate, one underexplored aspect is the association of certain protein families with tumor pathogenicity. We focused here on the G protein-coupled receptors family for three reasons. Firstly, it has been shown that this is the second most mutated class of proteins in cancer following kinases. Secondly, this family has been extensively studied resulting in a wide availability of experimental data for these proteins. Finally, more than 30 % of the drugs currently in the market target its members. For these receptors, we explored the mutational landscape across cancer patients compared to healthy individuals. Our findings show the existence of cancer-related alteration patterns that occur at conserved positions. Additionally, we computationally ranked these G protein-coupled receptors on their importance in the pathogenesis of cancer based on multiple objectives. The result is a list of recommendations on where to focus next. These results suggest that there is room for repurposing existing therapies for cancer treatment while also highlighting the risk of potential interactions between cancer treatments and common drugs. All in all, we present a window of opportunity for new targeting strategies in oncology for G protein-coupled receptors.

## Introduction

Cancer is the second leading cause of death globally [1]. Research towards this multifactorial disease has expanded our knowledge significantly over the last two decades [2,3]. Recently, results from these endeavors have been condensed in the form of public databases containing patient-derived data [4]. Cancer is typically the result of compounding mutations that transform healthy cells to malignant ones [5]. Previous work involving large scale mutational analysis picked up G Protein-coupled receptors (GPCRs) as the second most mutated class of proteins in the context of cancer [6]. Cancer cells are driven to proliferate and avoid the immune system. GPCRs have multiple functions in this process from increased growth (early stage) all the way to metastasis (late stage) [7]. Thus, any anomalies in GPCR functioning might be related to cancer growth. Another interesting property of GPCRs is that they are the most common drug target family with around 35% of drugs acting through a GPCR [8], providing a diverse set of molecular tools to potentially combat cancer.

GPCRs consist of seven highly conserved transmembrane (TM) domains, which often serve as a ligand binding pocket for their natural ligands, e.g. endogenous hormones or neurotransmitters. In addition, these TM domains are connected via extra- and intracellular loops (ECL; ICL) displaying a lower degree of conservation [9]. Most GPCRs also have an eighth TM domain that is connected by intracellular loop 4. The extracellular loops are known to also be involved in ligand recognition and activation, whereas the intracellular part of the receptor is linked to G protein recognition and activation. Finally GPCRs contain an N- and C-terminus which are also relatively little conserved [10,11]. To denote the residues in GPCRs in a comprehensive way, we use Ballesteros-Weinstein (BW) numbering [12]. BW is mainly restricted to the TM domains and consists of two parts, i.e. the first number is the TM where this residue is found, and the second number is relative to the most conserved residue in that TM. The most conserved residue receives number 50, and the number goes down for residues towards the N-terminus and up for residues towards C-terminus.

In previous work, knock-down studies have been performed on several proteins to identify their role in the context of cancer, but these studies were typically embarked upon after prior identification of the protein’s role in cancer [13–15]. One of the main reasons these *in vivo* studies are done is to identify whether a mutation is either a driver or a passenger mutation, where the first provides a selective growth advantage, and thus promotes cancer development, while the latter has occurred coincidentally and is thus generally of less interest. Moreover, these studies provide insight whether a driver mutation is located on either an oncogene or a tumor suppressor gene [16].

In the current work, we focused on GPCRs in the context of cancer by using patient-derived data sets and specifically looked at trends and mutational patterns in this protein family. We performed a deeper investigation into several “motifs”, parts of the GPCR sequence that are conserved that contribute most to the stability and function of the GPCR [17–19]. Moreover, we provide a list of GPCRs with known small molecule ligands (in some cases approved drugs), ranked by relative interest for follow-up using multi-objective ranking. This ranking incorporates mutational count, locations of mutations in regions of interest, availability of in-house expertise, and ability to perform virtual screening (as performed by QSAR). Finally, we exemplified our findings in a more in-depth analysis on C-C chemokine receptor type 5 (*CCR5*) to show the feasibility of our approach.

## Results

### Overview of datasets

Several filtering steps were applied to both the GDC and 1000 Genomes dataset including constraints to missense mutations and to mutations in GPCRs in residues defined in the GPCRdb alignment. The mutation analysis was done for all unique missense mutations in GPCRs, while for the Pareto optimization the datasets were enriched with ChEMBL data for those GPCRs for which such data were available. The corresponding numbers are shown in Table 1.

**Table 1:**
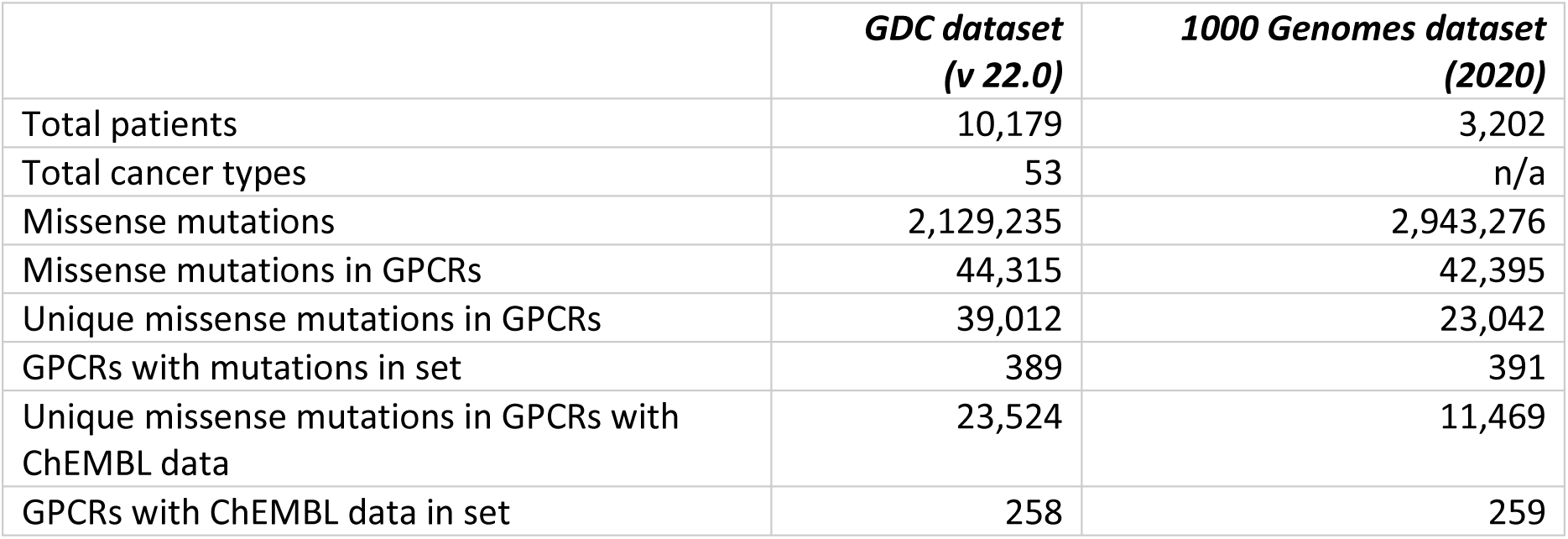
Overview of the composition of the GDC and 1000 Genomes datasets.

The GDC dataset is larger compared to the 1000 Genomes set based on patient count, but both are in the same order of magnitude when looking at the amount of missense mutations. Nevertheless, for better comparison in our analyses, the fraction of mutated residues per dataset was used instead of absolute mutation count to correct for the absolute difference in data points.

### Two-Entropy Analysis

A two-entropy analysis (TEA) was performed on our dataset as was done previously with slight modifications [19,20]. Key to the TEA approach is that for each alignment position the Shannon entropy is calculated both within a GPCR subfamily and within all GPCRs, with the difference between these indicating the residue function in the protein family and superfamily. From this type of analysis multiple interesting groups were identified, in particular residues relevant for receptor function such as activation (type A), and residues relevant for ligand recognition (type L). The former is made up by positions with a low Shannon entropy both within GPCR subfamilies and for the entire GPCR superfamily, indicating high conservation in general and within the subfamily. This high conservation has been linked to their involvement in GPCR-conserved working mechanisms [20]. The latter (type L) is made up by residues that are conserved within subfamilies, yet are not so much conserved within the GPCR superfamily. Hence, these are typically associated with ligand recognition, which is specific and conserved within a given subfamily. Type L residues are represented in the top left corner in Figure 1, but in our analysis this trend is less clear. This is likely as we have not limited ourselves to one family such as Class-A GPCRs (thus increasing the overall entropy). Despite the shift of type L positions, the positions from the original TEA analysis end up in the expected location. Moreover, in the top right corner of Figure 1, a third group of residues is represented: those that are conserved neither among all GPCRs nor GPCR subfamilies. These are more likely to have only a small implication in relevant receptor functions.

**Fig 1:**
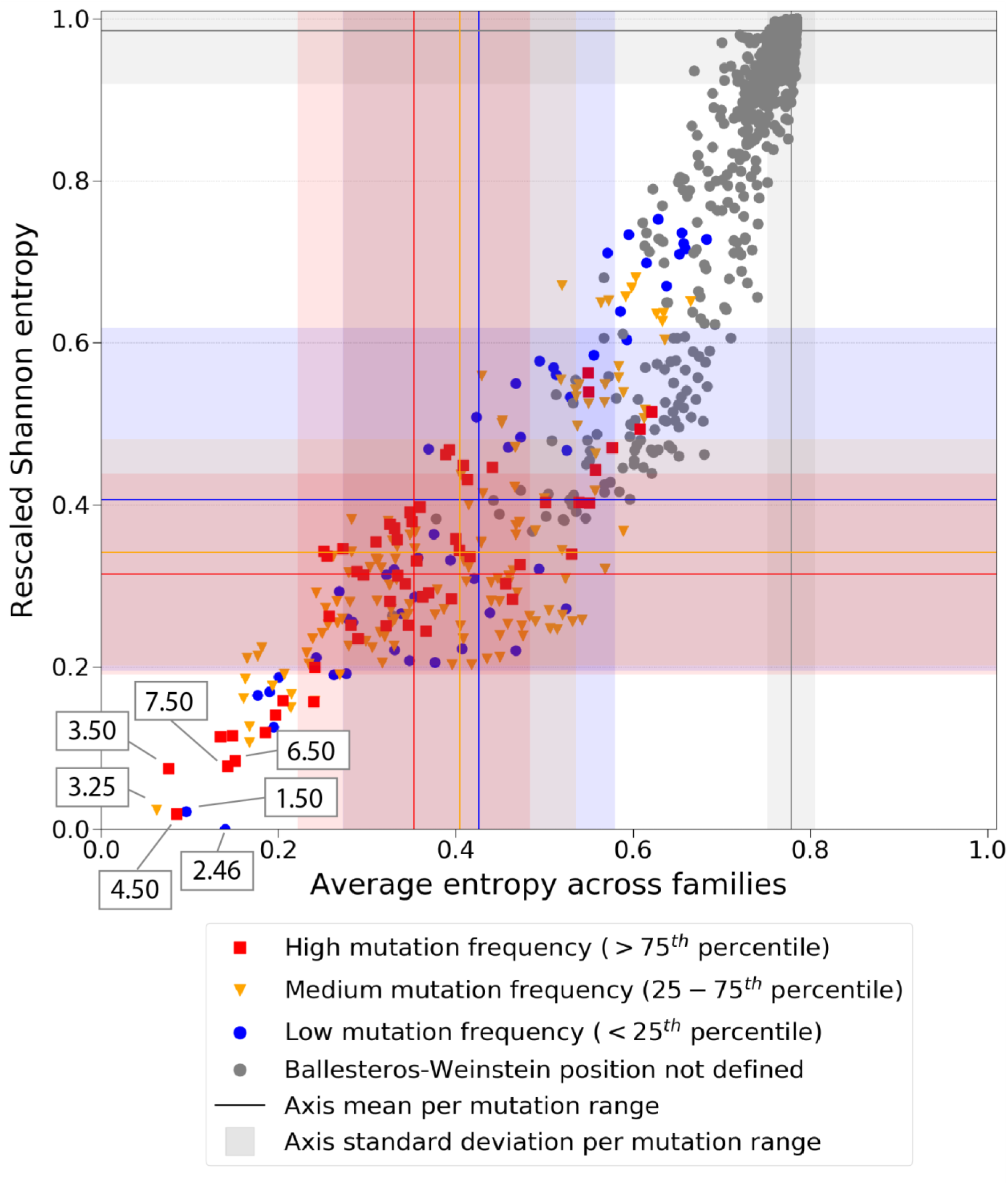
Shannon entropy across GPCR families versus Shannon global Entropy correlated to cancer-related mutations. A two-entropy analysis plot for all GPCRs with aligned positions. The average entropy across families, i.e. conserved within a family is on the x-axis, and the Shannon entropy overall on the y-axis. Residues are colored by the frequency of mutations found in the GDC dataset, with blue being low (< 25^th^ percentile), orange medium (25-75^th^ percentiles) and red high (> 75^th^ percentile). Residues with no defined Ballesteros-Weinstein labels are colored grey. Blue, orange, red, and grey lines represent the mean entropy values for each axis per mutation range (high, medium, low, and non-defined Ballesteros-Weinstein, respectively). Blue, orange, red, and grey shadows represent the standard deviation to the mean entropy values for each axis per mutation range (high, medium, low, and non-defined Ballesteros-Weinstein, respectively).

In Figure 1, we colored the data points based on the frequency of absolute mutation counts found per Ballesteros-Weinstein GPCR aligned position in cancer patients in the GDC dataset. The rest of aligned positions without a Ballesteros-Weinstein label is represented in Figure S1. We defined residues with a high mutation frequency as those above the 75^th^ percentile in the distribution of mutation counts by position. Conversely, residues with a low mutation frequency were defined as those under the 25^th^ percentile. The middle mutation frequency are the remaining data points. From Figure 1, it follows that absolute mutation count is (anti)correlated with entropy. We observe a trend where the more conserved type A residues (bottom left corner of the graph, low entropy) have a higher mutation rate in cancer compared to the less conserved residues (top right corner of the graph, high entropy). We illustrate this with the mean ± SD entropy overall and across families, represented in Figure 1 for each mutation range. The low mutation range has mean entropy values of 0.41 ± 0.21 and 0.43 ± 0.15 (Shannon and Average entropy across families, respectively). Meanwhile, the high mutation range has lower mean entropy values of 0.31 ± 0.12 and 0.35 ± 0.13, respectively. On the contrary, the trend is not observed on natural variance data from the 1000 Genomes dataset (Figure S2). There, the mean entropy values for the low mutation range are 0.39 ± 0.20 and 0.41 ± 0.16, respectively; and 0.32 ± 0.01 and 0.42 ± 0.11 respectively for the high mutation range. Comparing the GDC data and 1000 genomes data we observe an average downward shift in entropy values for highly mutated positions per subfamily (not in the overall Shannon entropy) and an upward shift for low mutated positions. Combined this shows a pressure in the GDC data for mutations in subfamily specific positions at the expense of mutations in non-conserved positions. This trend is supported by the fact that from the type A residues highlighted in Figure 1, the higher mutation frequencies are associated with the most conserved positions in TM domains 3, 4, 6, and 7 (i.e. 3.50, 4.50, 6.50, and 7.50). Three of these (i.e. 3.50, 6.50, and 7.50) are interesting positions, since they are part of the “DRY” (TM3), “CWxP” (TM6), and “NPxxY” (TM7) conserved GPCR functional motifs. The high amount of mutations in residues of these motifs is remarkable and will be investigated further in section ‘*Mutation patterns within functionally conserved motifs’.* Overall, cancer mutation frequency is correlated with individual residue conservation as we initially noted from Figure 1. We therefore investigated groups of residues as defined by GPCR domains in order to further explore cancer mutation patterns.

### Mutation rates over GPCR domains

We hypothesized that mutations associated with altered function in the context of cancer, would occur more frequently in the domains with higher conservation (i.e. TM domains) where positive selective pressure would favor those that alter receptor function. Conversely, we would expect mutations to be distributed more randomly over the sequence among the 1000 Genomes set and to be underrepresented in the conserved TM domains. However, the distribution in both sets appears to be quite similar (Figure 2A). Looking at the absolute counts per protein domain, we observe that most mutations are in the N-terminus (∼ 25% of the total), followed by the C-terminus (∼ 15% of the total), which are on average the longest domains compared to loops and TM domains. Around 40% of the mutations are found in the aggregated 7TM, but individual transmembrane domains follow in mutation count the N- and C-terminus together with ICL3 and ECL2, with the remainder of the loops having the lowest amount of mutations. The cause for this is most likely twofold; on the one hand both the N- and C-terminus are not as conserved as the 7TM domains of the GPCR, on the other hand the domain lengths are higher and hence the chance of mutations occurring is equally higher. Comparing the mutation fractions (percentage of absolute mutations found per domain) across the different domains shows no major differences between GDC and 1000 Genomes derived data, although there is enrichment observed in cancer-related mutations in the TM regions, as opposed to what is observed for the N-terminus and C-terminus. However, from these data it is difficult to conclude that cancer mutations have a specific domain they aggregate on. In order to remove the bias mentioned above regarding average length of the different domains, we corrected for average length of the domains by dividing the total amount of mutations per domain in the data set by the average length of this domain resulting in the average amount of mutations per position. Subsequently, these values were scaled between zero and one for better comparison with the newly calculated property: Rescaled average number of mutations per position over domains (Figure 2B). Both the absolute counts in the domains and an aggregated overview are represented in Figure S3.

**Fig 2:**
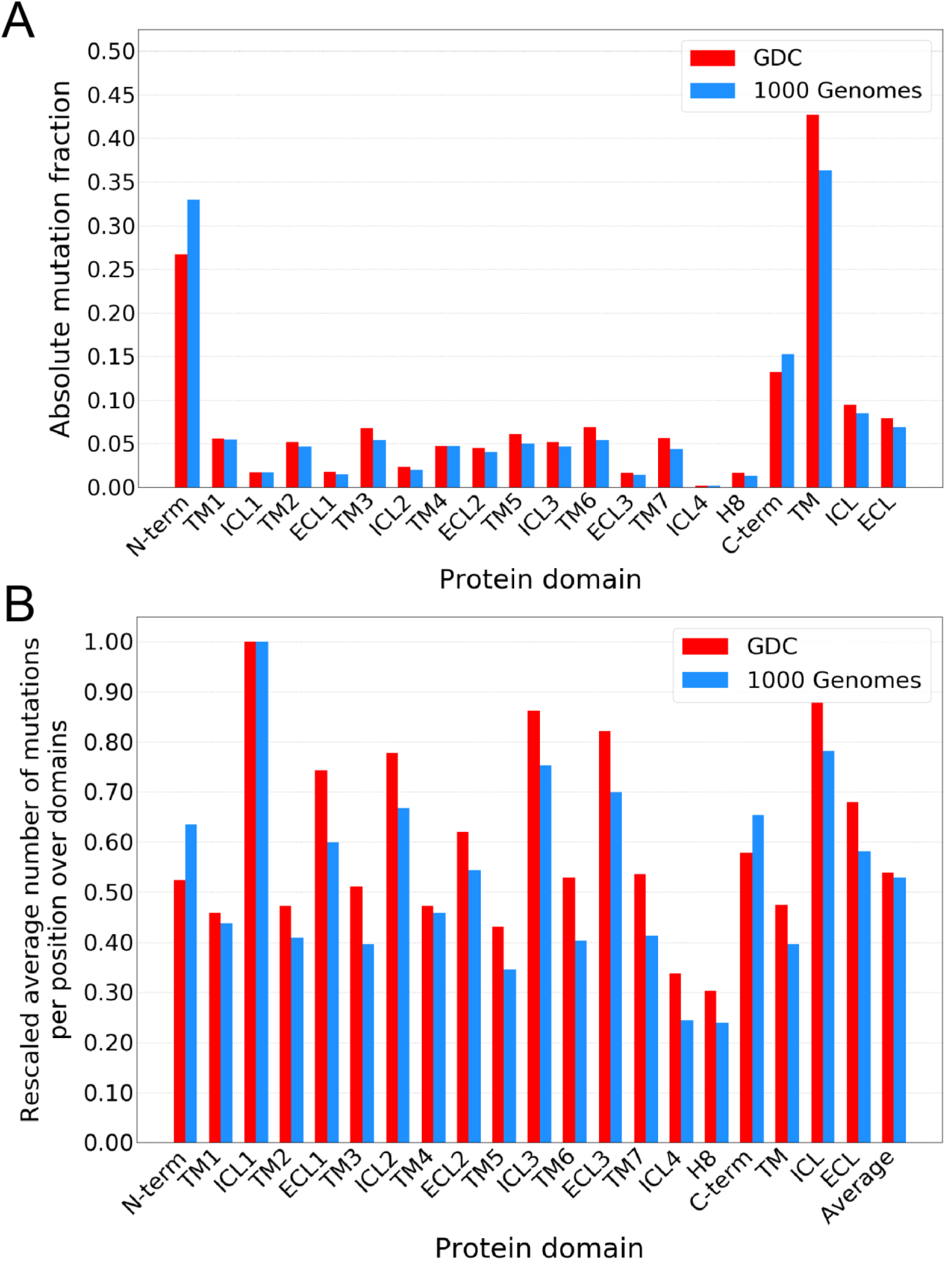
Mutation fractions per GPCR protein domain. (A) Mutation fraction from the total number of mutations found in the GDC and 1000 Genomes data, split per GPCR domain that they were found in. (B) Mutation count corrected for protein domain length and scaled between zero and one for better comparison. Scaling was done between absolute zero and absolute maximum for visualization purposes. “TM”, “ICL” and “ECL” represent the aggregated rescaled average number of mutations per position for those domains, and ”Average” is the average rate over all the data. Red bars shows the mutation rates in the GDC dataset, while blue bars shows the rates of the 1000 Genomes dataset.

The rescaled average number of mutations per position over domains represented in Figure 2B allow us to more accurately describe the differences between protein domains and datasets. In this figure, the mutation rates on average over the whole GDC and 1000 Genomes datasets are also shown for reference. For the highest scoring domain, ICL1, 656 mutations were found in the GDC dataset. This domain has an average length of 4.54 residues, resulting in 144.5 mutations per residue, which was then scaled to a score of 1.00. This domain was also the highest scoring in the 1000 Genomes dataset. These values represent almost double the average rate over the whole data, which is 0.54 and 0.53 for the GDC and the 1000 Genomes datasets, respectively. Conversely, H8 has by far the lowest rate of mutations, which might be due to this domain, previously associated with mechanosensitivity of GPCRs, not being present in all receptors [21]. The length of ICL and ECL loops, logically, is considerably more variable than that of TM domains and, in the case of ICL1, the higher mutation rates are not necessarily found in the most conserved alignment positions (i.e. 12.48-12.50). These observations make us suggest that the mutations in ICL1 are context-specific and need to be examined for every GPCR individually. However, the effects of these mutations may be limited [22–24]. We observed that all TM domains, all ECL loops, and ICL2-4 demonstrate a slightly higher mutation rate in the cancer samples, although for TM1 and TM4 the difference is minimal. Conversely, N-terminus, ICL1, and C-terminus demonstrate a slightly lower mutation rate in the cancer samples. From the analysis of this data, we concluded that some domains may be more amenable to mutation in the context of cancer, but that the high diversity of the GPCRs studied and their diverse roles obscure a clear conclusion. To further investigate these incipient mutation patterns in protein domains, we proceeded to the analysis of previously identified motifs that have a conserved function in GPCRs and that were also highlighted in our two-entropy analysis.

### Mutation patterns within functionally conserved motifs

Several highly conserved motifs relevant for GPCR function are known, in which amino acid point mutations usually cause a disruption or change in function [25–30]. The “DRY” motif is important for receptor activation, whereas both the “DRY” and “NPxxY” motifs were found to be instrumental in stabilization of the receptor-ligand complex, contributing a significant portion to the stability of the helices. Finally, the “CWxP” motif is important for receptor activation as it enables movement of the helices [26,27,31]. To be able to compare these motifs, which are of different lengths, we calculated an average mutation rate for each, correcting for the length difference, similar to Figure 2B, and with a comparable scaling from zero to one. As a reference, the average mutation rates obtained over the whole GDC and 1000 Genomes datasets are also shown.

From Figure 3 it follows that for each motif and its six neighboring residues (deemed extended motif), there is an increase in mutations in cancer patients compared to the natural variation. At the same time, there is a similar average rate of mutations per residue in the GDC dataset and in the 1000 Genomes dataset (column Average in Figure 3). Moreover, in the GDC dataset (red bars) “DRY” is enriched for mutations in samples collected from cancer patients compared to the average mutation rate whereas for the 1000 Genomes (blue bars) there is a clear reduction in mutation rate visible. In the extended “DRY” motif, this effect is smaller but still visible. For both “CWxP” and “NPxxY” the rate in the GDC dataset is comparable to the whole sequence rate, whereas it is also lower in the 1000 Genomes dataset for these two motifs. In the extended motifs of “CWxP” and “NPxxY”, this trend is still observed. An absolute count of the mutations found per residue in the aforementioned motifs in both the GDC dataset and the 1000 Genomes dataset is shown in Figure S4. From this, we conclude that a trend of higher mutation rates is present within highly conserved motifs in the GDC dataset compared to the average mutation rate, which is not observed in the 1000 Genomes dataset. Moreover, a pattern is observed that the mutation rate in conserved motifs is lower in the 1000 Genomes set compared to the GDC set, confirming their essential role and conservation. To gain further insights into the mutation rate and trends identified here we selected the most mutated individual positions in the GDC dataset (Figure 4). A count overview of unique GPCR cancer mutations for Ballesteros-Weinstein positions is provided in Figure S5, and an overview of the substitutions found in all of the mutations is provided in Figure S6.

**Fig 3:**
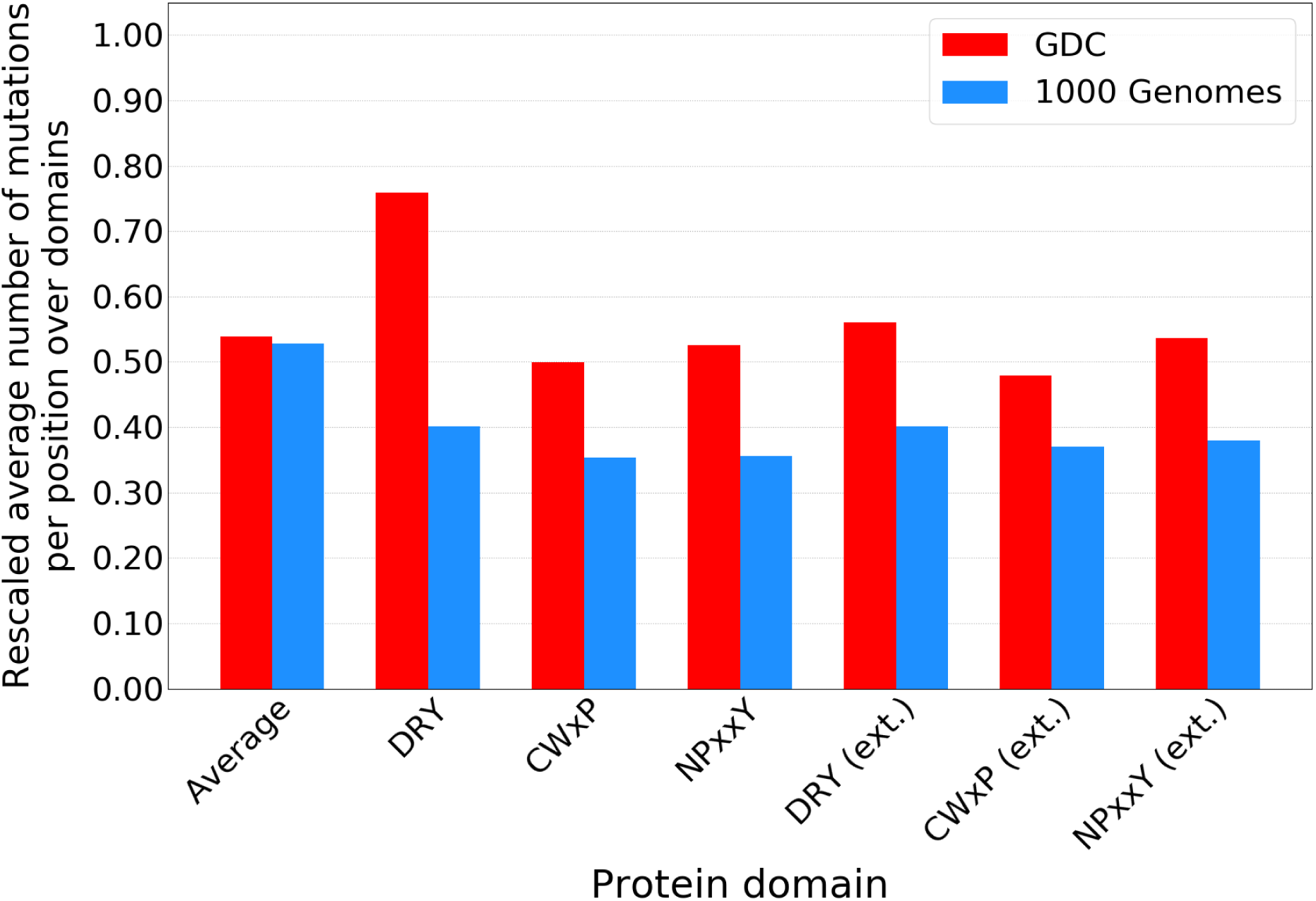
Mutation fraction of conserved motifs and their surrounding residues (extended). Mutation rates in GDC and 1000 Genomes datasets of conserved motifs found in GPCRs. “DRY”, “CWxP”, and “NPxxY” motifs are analyzed along with their “extended” version, which includes three residues before and after the motif, as found in Table 2. “Average” is the average rate over all the data. Red bars show the mutation rate in the GDC dataset, while blue bars show the rate of the 1000 Genomes dataset.

**Fig 4:**
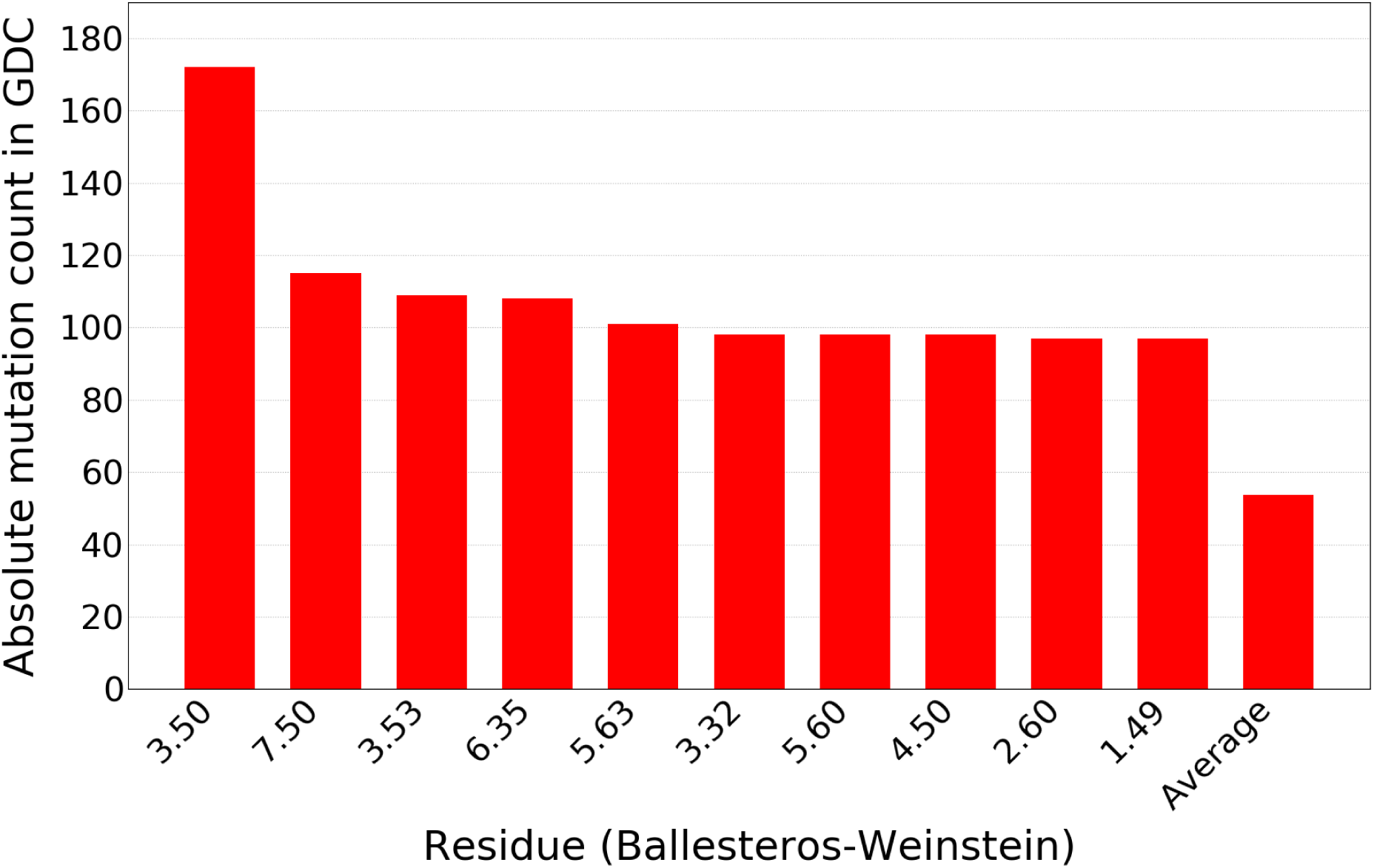
Most frequently mutated residues in GDC. The top 10 most frequently mutated positions found in GPCRs in the GDC dataset. The residue location in Ballesteros-Weinstein notation is shown on the x-axis, while on the y-axis the mutation count of that residue is given. “Average” is the average mutation count per residue over all the data.

In Figure 4 it is shown that the highly conserved positions in TM domains 3, 4, and 7 (3.50, 4.50 and 7.50) all appear in the top 10 of most frequently mutated residues. These TM domains already demonstrated a trend in the domain analysis and the locations are part of the “DRY” and “NPxxY” motifs. These three positions also showed a higher mutation rate in the two-entropy analysis. In addition, residue 3.53, which is part of the extended “DRY” motif, also shows up as highly mutated. The fact that two residues of the “DRY” motif are some of the most mutated in cancer could explain why this motif shows the biggest enrichment in the GDC dataset in Figure 3. On the contrary, no residues of the “CWxP” motif are included in Figure 4, which aligns with this motif showing the smallest enrichment in Figure 3. Disruptions in these motifs due to mutations can influence GPCR function in several ways, explaining the enrichment patterns in cancer patients compared to natural variance observed in our analysis.

### Ranking GPCRs for follow up

Having confirmed that patterns can be identified in GPCR mutations in the cancer context, we ranked GPCRs for experimental follow-up. For each GPCR, the absolute mutation count was divided by receptor length, to provide a mutation rate for each receptor (a higher mutation rate yielding a lower – better – rank). To identify patterns within GPCR families (as classified by the GPCRdb [32]), a family-wide rank was calculated by averaging the ranking of each of the members in a family and subsequently compared to the other families (top 20 shown in Figure 5).

**Fig 5:**
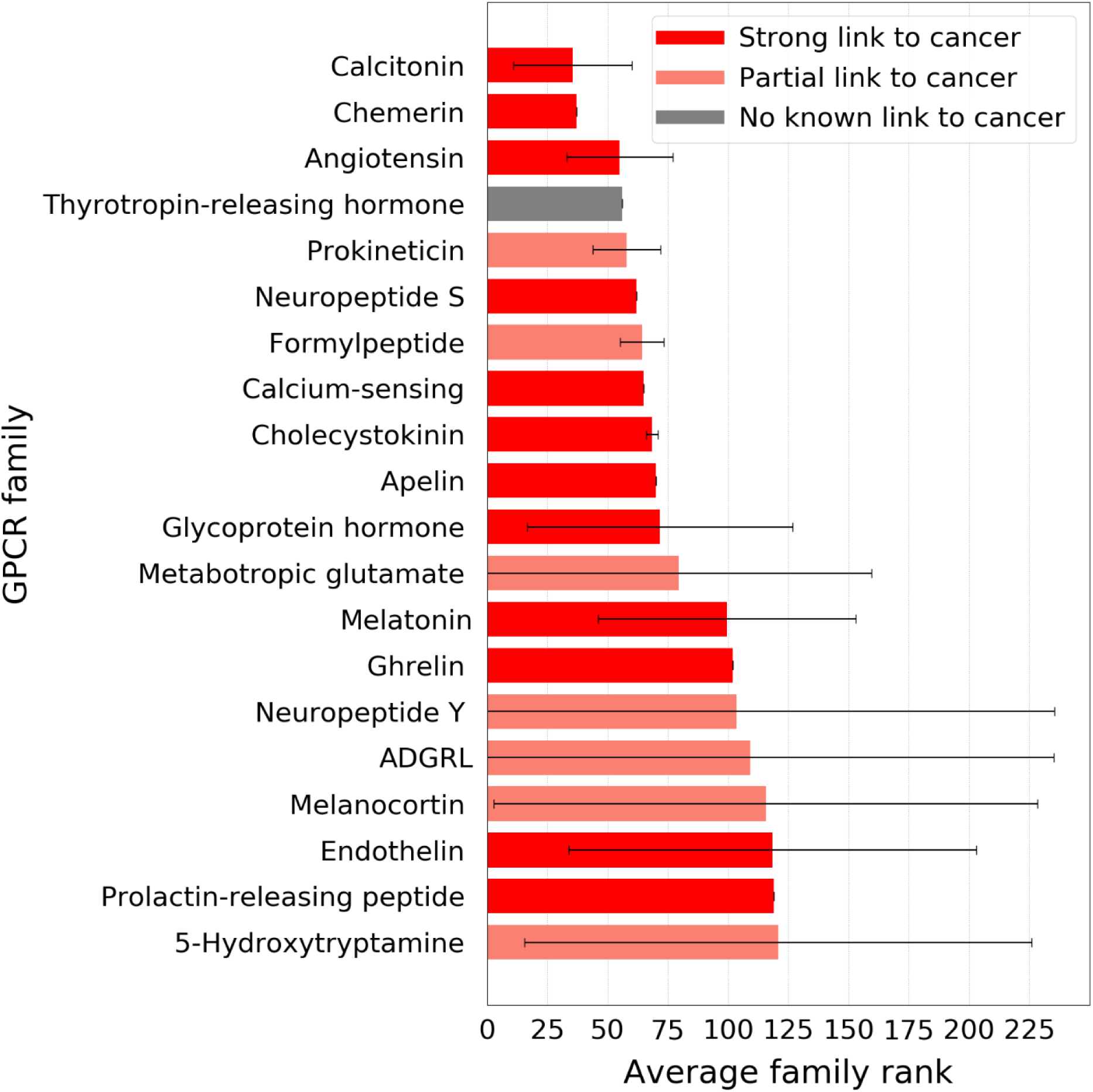
Average Rank of GPCR families and their link to cancer in the literature. Average rank of GPCR families related to the mutation rate in individual family members. Shown on the y-axis are the different GPCR families as categorized by GPCRdb, while on the x-axis their average rank as a receptor family is given. The lower average rank value, the better. The error bars represent standard deviation of individual GPCR rankings within the family. Color-coding represent the link to cancer in the literature for the family. Red represents a strong link (i.e. all members of the family have been linked to cancer), salmon represents partial link (i.e. some members of the family have been linked to cancer), and grey represents no link to cancer reported.

The majority of GPCR families identified in the top 20 ranking have been previously linked to cancer in the literature, as the color-coding in Figure 5 represents. For most of these families, such as the calcitonin, the angiotensin, or the melatonin receptor families, all individual members of the family have been linked to cancer (red bars in Figure 5) [33–38]. For some others, however, only some members of the family have been associated with cancer in the literature (salmon bars in Figure 5), such as the metabotropic glutamate (mGlu) or 5-hydroxytryptamine (serotonin) receptor families [39,40]. Notably, the family with the fourth best ranking (i.e. thyrotropin-releasing hormone receptor family), has not been previously linked to cancer in the literature yet the ligand for this family has been linked to cancer and cancer-related fatigue [41,42]. The standard deviation represented by error bars in Figure 5 gives an idea of the differences in individual member rankings within families. Families with a standard deviation of zero correspond to families with only one member, since every receptor has a unique mutation rate and rank in this set. Of note, families with a higher standard deviation correspond to those that have been partially linked to cancer, with a few exceptions (i.e. glycoprotein hormone receptor and endothelin receptor families). Hence, big error bars represent ranking differences of the family members. The retrieval of families previously identified in the context of cancer validates our approach and opens room for further target selection strategies based on mutagenesis data in cancer.

To further narrow the list of selected receptors, Pareto sorting was performed to identify GPCRs with a suggested high impact in cancer biology that may be amenable to small molecule intervention and follow up study. Pareto sorting is a means to sort a list of items based on multiple (not always correlating) properties. The feasibility of small molecule intervention was assessed by training a machine-learning model (random forest [43]) for each GPCR in our data set using bioactivity data from ChEMBL 27 [44,45], with circular fingerprints as molecular descriptors [46]. The selected properties for Pareto ranking were: Mutation rate in TM domains in GDC (maximized), mutation rate in TM domains in 1000 Genomes (minimized), average R^2^ of ChEMBL QSAR prediction models (maximized), and in-house availability of proteins for experiments (maximized). The order of the properties determine the priority during the Pareto sorting.

The first front in the Pareto optimization is considered “dominating”, which means that this set of GPCRs has no GPCR that scores better in the properties. For the remaining (i.e. dominated) data points, a second front can be calculated, with GPCRs that score worse than those in the first front but better than the rest of the solutions. Therefore, we used the first and second fronts for a subsequent ranking based on crowding distances between the receptors (Figures 6A and 6B, respectively). Crowding distances are a measure of how dense the environment is. Denser environments mean more balance in the objectives and thus more interesting GPCRs. As the crowding distance can go up to near infinite, we used a cut-off at a value of 10.

**Fig 6:**
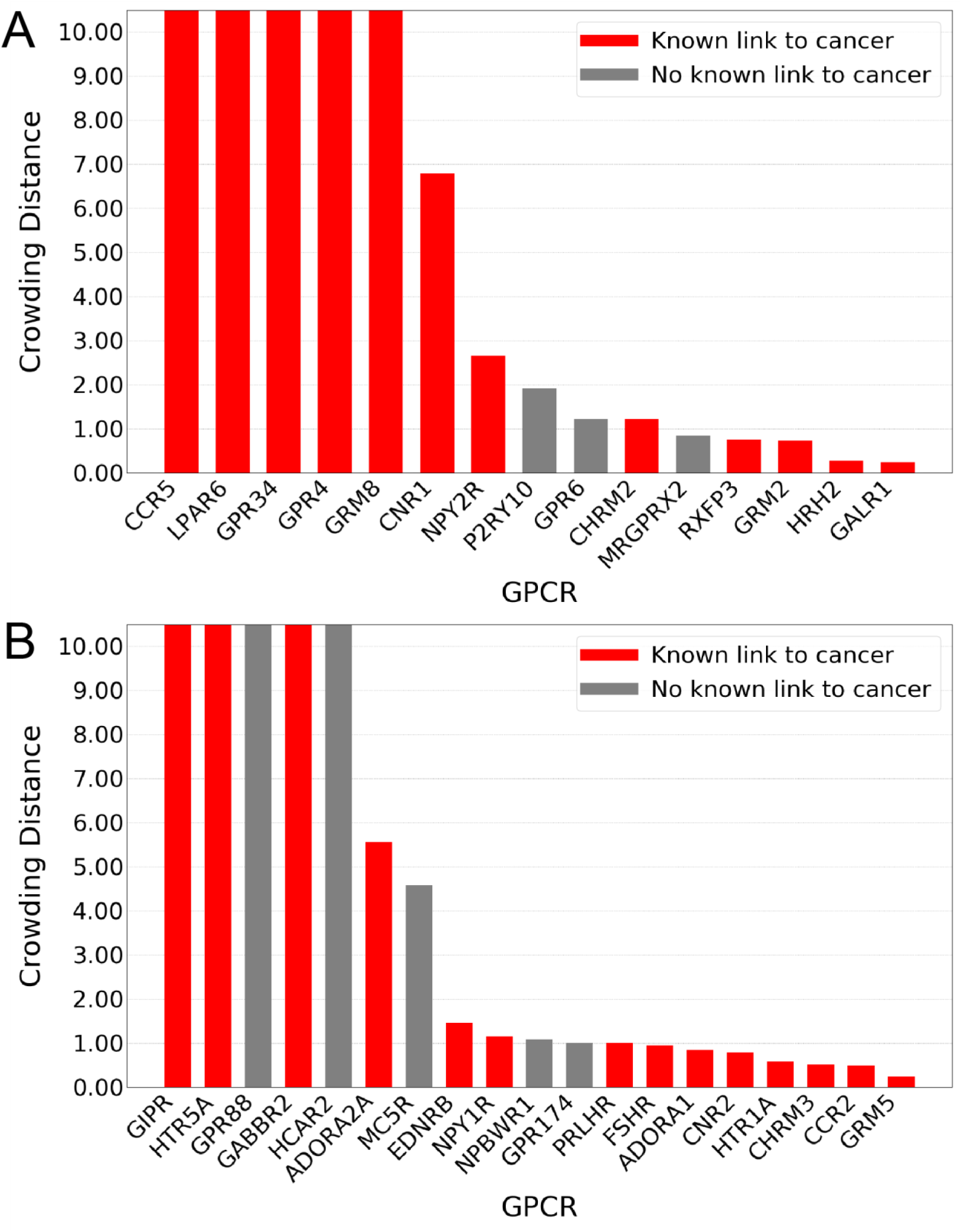
Crowding distances of the first and second Pareto fronts. (A) First Pareto front, consisting of 15 GPCRs. (B) Second Pareto front, consisting of 19 GPCRs. On the x-axis the gene names of GPCRs are shown, while on the y-axis their crowding distance is shown. Crowding distance was cut-off at 10, as the differences between these high-scoring receptors become negligible above that threshold.

In Figure 6A, the 15 GPCRs from the best scoring (first) front are shown, which translate to the GPCRs with the most desirable scores in the combined objectives of the Pareto optimization. We demonstrate that GPCRs previously linked to cancer show up in the first front alongside others that have not been thoroughly investigated yet. Hence, our list can be seen as a list of potential candidates for follow-up experimental research. Twelve of these GPCRs (80%) have been identified in literature as related to cancer (red bars in Figure 6A). The second Pareto front (Figure 6B), reveals a list of 19 GPCRs, from which 14 (74%) have been previously linked to cancer (red bars in Figure 6B). Out of the cancer-related receptors in our analysis we selected one of the top entries of our first Pareto front, *CCR5*, as a case study for further investigation and performed a structural analysis based on its crystal structure to characterize the potential effects of the retrieved mutations in receptor function and/or ligand binding.

### CCR5 structural analysis

Mutations found in the GDC dataset for *CCR5* were cross-linked to GPCRdb data, to find previously published mutagenesis data. We then mapped the mutations on a 3D crystal structure of the receptor (PDB code 4MBS [47]). In this structural analysis, we focused on regions relevant for protein function and ligand binding. As shown in Figure 7A, these mutations are widely spread across the receptor’s structure, including mutations in ECL2 - a region that largely contributes to chemokine ligand recognition (Figure 7B) [48], G protein binding region, and orthosteric binding site (Figures 7C and 7D). The crystal structure of *CCR5* used as a reference in Figure 7 (PDB code 4MBS) contains the thermostabilizing mutation *A233^6.33^E*, which has been characterized for the inactive *CCR5* conformation. In this structure, a small molecule inhibitor - maraviroc - is co-crystalized in the orthosteric binding site (i.e. spanning the so-called major and minor binding pocket [49]), as shown in Figure 7D. Of note, some of the mutations found in the GDC dataset are in positions in close proximity to the inhibitor. Out of the 73 mutations found in our dataset, only 12 mutations had been previously annotated, while 37 mutations had no data available and 24 consisted of not-annotated data. Further analysis of previously annotated data shed some light on the functional implications of these mutations, as discussed below.

**Fig 7:**
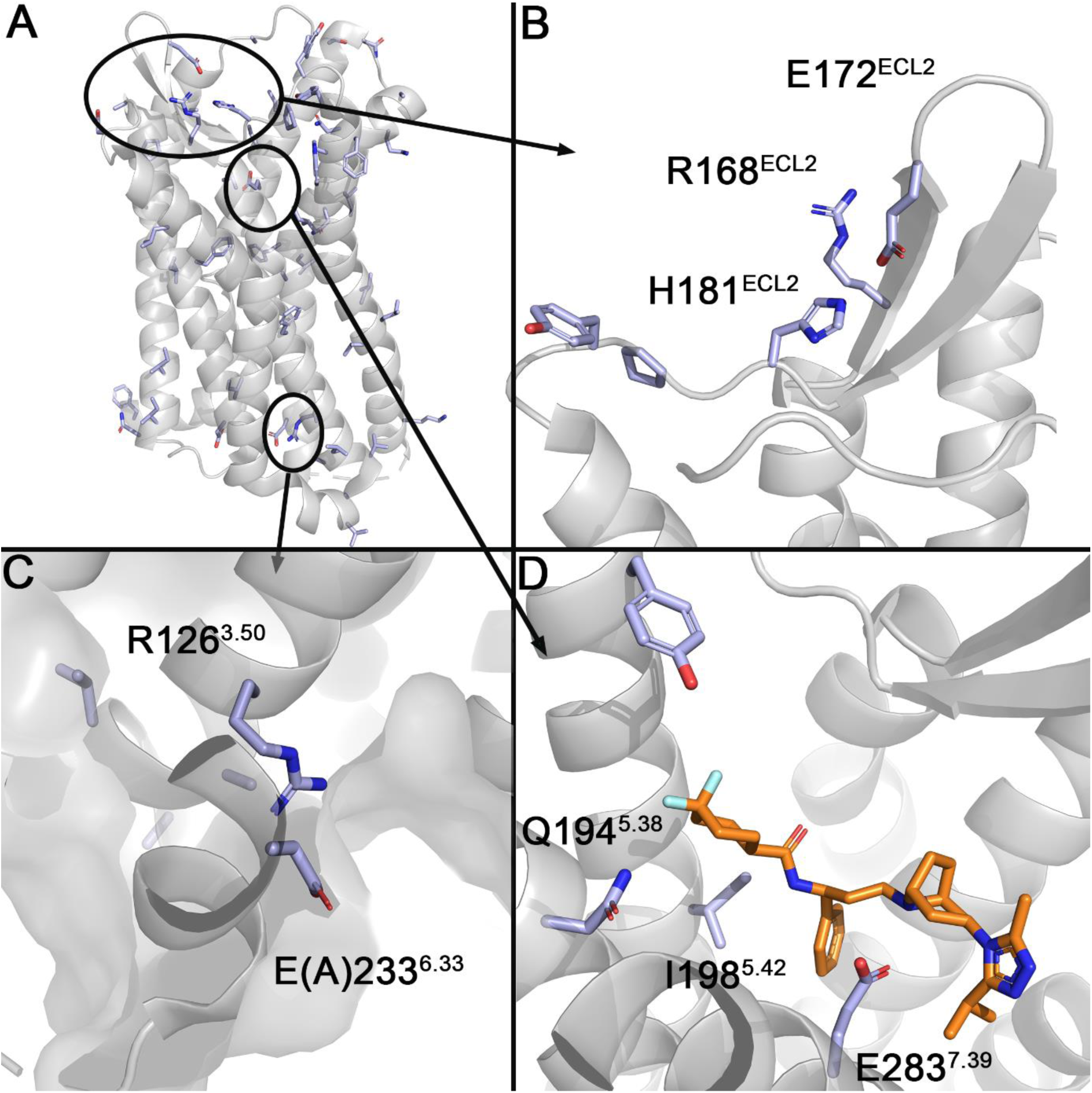
Cancer-derived mutation mapping in CCR5 structure. (A) The mutations found in the GDC dataset for CCR5 mapped on the 3D structure of the receptor. (B) Mutated residues found in ECL2 region. (C) G protein binding site, containing the mutation A233^6.33^E, which has been characterized as a thermostabilizing mutation for the inactive CCR5 structure (PDB code 4MBS). (D) The orthosteric binding site, with the small molecule inhibitor maraviroc (orange).

## Discussion

In this study, we performed a comprehensive comparison of mutations found in cancer patients (GDC dataset) versus mutations found in natural variance (1000 Genomes dataset) in GPCRs. We followed this up by investigating several highly conserved motifs for an increase in mutation rate compared to the other residues. Finally, we performed a Pareto Front analysis to create a ranking of GPCRs that warrant follow-up for their context in cancer, and we analyzed some of the cancer-related mutations found for one of the top-ranking receptors from a functional-structural point of view.

Our original hypothesis was that more conserved residues (i.e. lower entropy in a two-entropy analysis of all residue positions in the GPCRdb alignment) would experience a higher mutational pressure in cancer patients. We confirmed this trend showing that positions with a low amount of mutations per position were assigned higher entropy values (0.41 ± 0.21 and 0.43 ± 0.15 Shannon and Average entropy across families, respectively, as shown in Figure 1) than positions with a high amount of mutations per position (0.31 ± 0.12 and 0.35 ± 0.13, respectively). Conversely, the trend was not observed in a similar analysis correlated to mutations in the 1000 Genomes dataset (Figure S2). In the original two-entropy analysis by Ye *et al.,* focused on class A GPCRs [20], the algorithm enabled the identification of residues involved in ligand recognition, but that trend was not as clear in our analysis. These observations implied a decreased advantage of the two-entropy analysis in an all-class GPCR analysis versus a class-A-only analysis. Overall, we identified an incipient pattern between evolutionary conservation and mutation rates in the GDC set, although this trend does not extend to the bulk of residues with intermediate entropies.

We also studied mutation distribution after aggregating residues by protein domains rather than exploring individual residues (Figure 2). Even though there were more mutations found in the larger domains such as the C- and N-terminus, when corrected for average length most of them showed similar mutation rates. Of note, mutations in TM, ICL, and ECL domains combined showed an enrichment in cancer patients versus natural variance, while the contrary was observed for the C- and N-terminus. Although the latter domains have been extensively described to play a role in receptor stabilization, signal transmission, and ligand and G protein recognition [50,51], they also represent the most variable domains in GPCRs in terms of length and motif composition, which explained the lack of an enrichment in cancer in these domains [52]. This aligned with the observation that GPCR mutation rates were not homogeneously distributed among cancer types, with some primary sites (e.g. Corpus uteri) showing a clear enrichment compared to others (see Figure S7 for more information). In literature the same has been suggested, with the emphasis on specific residue changes that affect the entire function of the protein [53–55].

A closer look at the more functionally conserved motifs of GPCRs showed a clearer pattern. The higher mutation pressure observed in the GDC data compared to the 1000 Genomes data to avoid these motifs, and especially the “DRY” motif (Figure 3), drove us to speculate that changes in these positions have a very high chance of disabling receptor function. Thus, this might not be tolerated in healthy tissues but can be advantageous to cancer development. For “DRY” mutations, it has been shown that G protein coupling and recognition can be decreased which can reduce binding affinity of drugs [17,29,31,56]. For both mutations in “DRY” and “NPxxY” it has been shown that a decrease in ligand-receptor complex stability may occur, decreasing the response from the GPCR [18,27,28]. Thus, any mutations found in these motifs can have an impact on the signal transduction of an endogenous ligand or the therapeutic effect of a small molecule drug. In fact, these motifs have been shown to be collectively involved in a conserved Class A GPCR activation pathway [57]. In practice, however, the effects of mutations found in these motifs have been shown to cause instability or loss of function in some GPCRs, but increased expression or activity in others [17,24,26,31,56,58–60].

Subsequently, we ranked in a multi-objective manner, via Pareto front analysis, the individual GPCRs for follow up work. In our ranking provided (Figure 6), approximately 80% of the top ranked receptors had a known link to cancer. Notable entries that have reported connections to cancer include the C-C Chemokine receptor (CCR) type 5, which has been linked to regulatory T cells mediating tumor growth [61], and type 2 as a key player in microenvironment-derived tumor progression [62,63]; LPA (Lysophosphatidic acid) receptor *LPAR6*, upregulated in bladder cancer [64]; GRM (Metabotropic glutamate) receptors 2 (*GRM2*) and 8 (*GRM8*), respectively known for dysregulating signaling pathways that are crucial in cancer prevention and activating variants in squamous cell lung cancer [65–67]; serotonin receptors 5HT_1A_ (*HTR1A*) and 5HT_5A_ (*HTR5A)*, the former known to be involved in at least breast, ovarian and pancreatic cancer, and the latter recently linked to breast cancer [39,68]; and the adenosine A_1_ (*ADORA1*) and A_2A_ (*ADORA2A*) receptors, linked to the progression and metastasis of a variety of cancer types as well as immune escape and immunotherapy [69–72]. The P2Y receptor family member 10 (*P2RY10*) is an example of GPCR not previously linked directly to cancer, found in the first Pareto front in rank eight. *P2RY10* has been linked to chemotaxis via eosinophil degranulation, which could make it a potential target in cancer [73].

Finally, the structural analysis of site-mutagenesis data in one of the top receptors from the first Pareto front (*CCR5*) shed some light into the functional implication of some of the cancer-related mutations. These include a cluster of six residues in ECL2 found within the GDC dataset, from which four positions had been previously shown to influence chemokine binding when mutated to Ala [74–76]. In the G protein binding site, the class A highly conserved R126^3.50^ was found to be mutated. This residue is part of the DRY motif and, as highlighted in the two-entropy analysis, it is the most frequently mutated position in the GDC set, resulting in altered G protein coupling to the receptor in for instance the adenosine receptor family [77]. Some experimental evidence is available for *CCR5* as well, where mutation of this residue to Asn abolished G protein signaling [56,78]. In the orthosteric site, four amino acids have been previously investigated by a site-directed mutagenesis study by Garcia-Perez *et al.*, namely Y187^5.31^, I198^5.42^, N258^6.58^, and E283^7.39^ [76]. The effect of these mutations in the binding affinity of a small molecule (maraviroc) and endogenous CCL5 chemokine recognition was variable. The biggest effect in the decrease of maraviroc binding affinity was observed for residue E283^7.39^, either when mutated to Ala or to the more conservative Gln. The structural effect of I198^5.42^ and E283^7.39^ mutations in maraviroc binding can be derived from the crystal structure of *CCR5* with this negative allosteric modulator (PDB code 4MBS, see Figure 7C). Indeed, mutations on these two positions had an important effect on the ligand binding of two other HIV-1 drugs - vicriviroc and aplaviroc - and clinical candidates - TAK-779 and TAK-220 - in two different studies [79,80]. It was further shown that, whilst *E283^7.39^A* abolishes maraviroc binding, chemokine CCL5 binding is mildly (20-fold) affected [79]. On the contrary, *Y187^5.31^A* showed almost no effect in the binding affinity of maraviroc, while affecting chemokine recognition [76]. These observations exemplified the relevance of our method to prioritize cancer-related mutations in site-mutagenesis studies, where they can be linked to receptor activation, endogenous ligand recognition, and the recognition of small (drug-like) molecules.

Recently, in a complementary extensive study by Wu *et al* (2019) [81] the TCGA dataset was used to identify significantly mutated GPCRs in cancer. Compared to their study, we elaborate on our findings through a motif analysis of highly conserved residues in GPCRs, a link to positional entropy, and a link to structural information (i.e. analyzing the *CCR5* chemokine receptor). Moreover, in our analysis we included the availability of chemical tools to study the selected GPCRs, as exemplified by our QSAR models. Recently, we have published an analysis of another GPCR, the Adenosine A_2B_ receptor, for which cancer-related somatic mutations were prioritized based on a structural analysis as presented here [46]. There we used a yeast system to explore the effect said cancer-related mutations have on receptor function directly and found that there is a complex pattern of activation modulation (increase, decrease, or disable). Similar approaches could be used to experimentally validate the relevance in cancer of somatic mutations in GPCRs prioritized in this work.

While in this computational approach the focus was on GPCRs, other receptor families can be investigated in a similar manner provided that there is a suitable dataset. Notable examples include solute carriers, or receptor-tyrosine kinases. The objectives in the Pareto optimization can also be adapted, providing a modified way of scoring the receptors depending on the scope of the study. Notably, our analysis was done focusing on differences in missense mutations occurring in cancer patients and natural variance. Nevertheless, many other alterations (e.g. insertion/deletions, gene and protein expression levels) have been reported for GPCRs in the context of cancer [82,83], and complementary analyses could be executed focusing on these. Finally, this computational approach can become part of a targeted therapy pipeline, suggesting key locations for *in vitro* and *in vivo* cancer-associated studies.

## Conclusions

We conclude from our study that mutations found in GPCRs related to cancer are in general weakly correlated to specific domains in the protein or evolutional conservation. Rather we conclude that these are highly context dependent (cancer type, tissue type). However, we do demonstrate that there is a higher mutational pressure in conserved motifs (i.e. “DRY”, “CWxP”, and “NPxxY”) in cancer patients (as shown in the GDC set) compared to healthy individuals. We observe a correlation between our mutational analysis and empirical findings on the role of several receptors in the cancer process. Moreover, we show that the role and mechanism of specific mutations can be elucidated using structural analysis as an intermediate step towards experimental validation. Finally, we have provided a list of GPCRs that are amenable to experimental follow-up based on our analysis. The provided data may help in exploring new avenues in the design of cancer therapies, either by linking existing data to ligand binding and recognition, or the identification of potential new roles for residues not previously studied.

## Methods

### Cancer related mutations

Cancer associated mutations were obtained from the Genomic Data Commons (GDC), part of the US National Cancer Institute effort [84], with the dataset used in this study obtained from their version 22.0 released on January 16^th^ 2020. GDC contains multi-dimensional mapping of genomic changes in several cancer types, including the complete dataset from The Cancer Genomic Atlas project (TCGA) [85]. As a means to facilitate reproducible, version-consistent, big data cancer data analysis, we re-compiled part of the GDC database version 22.0 in a MySQL [86] format. For this re-compilation, data was obtained from the GDC API engine, as well as their data transfer tool, depending on the availability. Exclusively unrestricted-access data was compiled. The SQL database contains 19 tables distributed in eight different fields, connected by a complex network of primary (PK) and foreign (FK) keys built to optimize the storage space and query processing. All PKs are unique numerical values. Some data fields (i.e. gene expression data) contain analyzed data derived from GDC raw data files. A more extensive description of the database architecture, as well as the analyses performed and the end-to-end mapping strategy is available in the supplementary data. For this project, we used data on somatic missense mutations found in a diverse set of cancer types, to which we will refer as the “GDC” data set.

### Natural variation

As reference for the analysis we used the 1000 Genomes data [87], including an additional data set released in 2020 by the New York Genome Center (NYGC). This is a dataset containing the natural variation of mutations in the genome. The dataset used in this study was obtained from Uniprot variance database in October 2020 [88]. From this data, all somatic missense mutations were gathered. From the extensive variance dataset, only mutations found in the 1000 Genomes subset were kept, this way removing cancer derived mutations from COSMIC and known pathological mutations [89]. Here we referred to this dataset as “1000 Genomes”.

### Mutation dataset curation

Using the aforementioned GDC and Uniprot databases, two filtering steps were applied (Figure 8). The first step yielded the missense mutations for all receptor families. In the second step, we filtered for GPCRs and aggregated their mutation data, ending up with GPCR-unique mutation pairs, along with the frequency, while still being able to find single mutations. The second filtering step also annotated the resulting GDC and 1000 Genomes datasets with identifiers from GPCRdb [90]. In a later step, prior to QSAR modelling and Pareto sorting, both datasets were enriched with bioactivity data from ChEMBL (release 27) [45,91] (see below).

**Fig 8:**
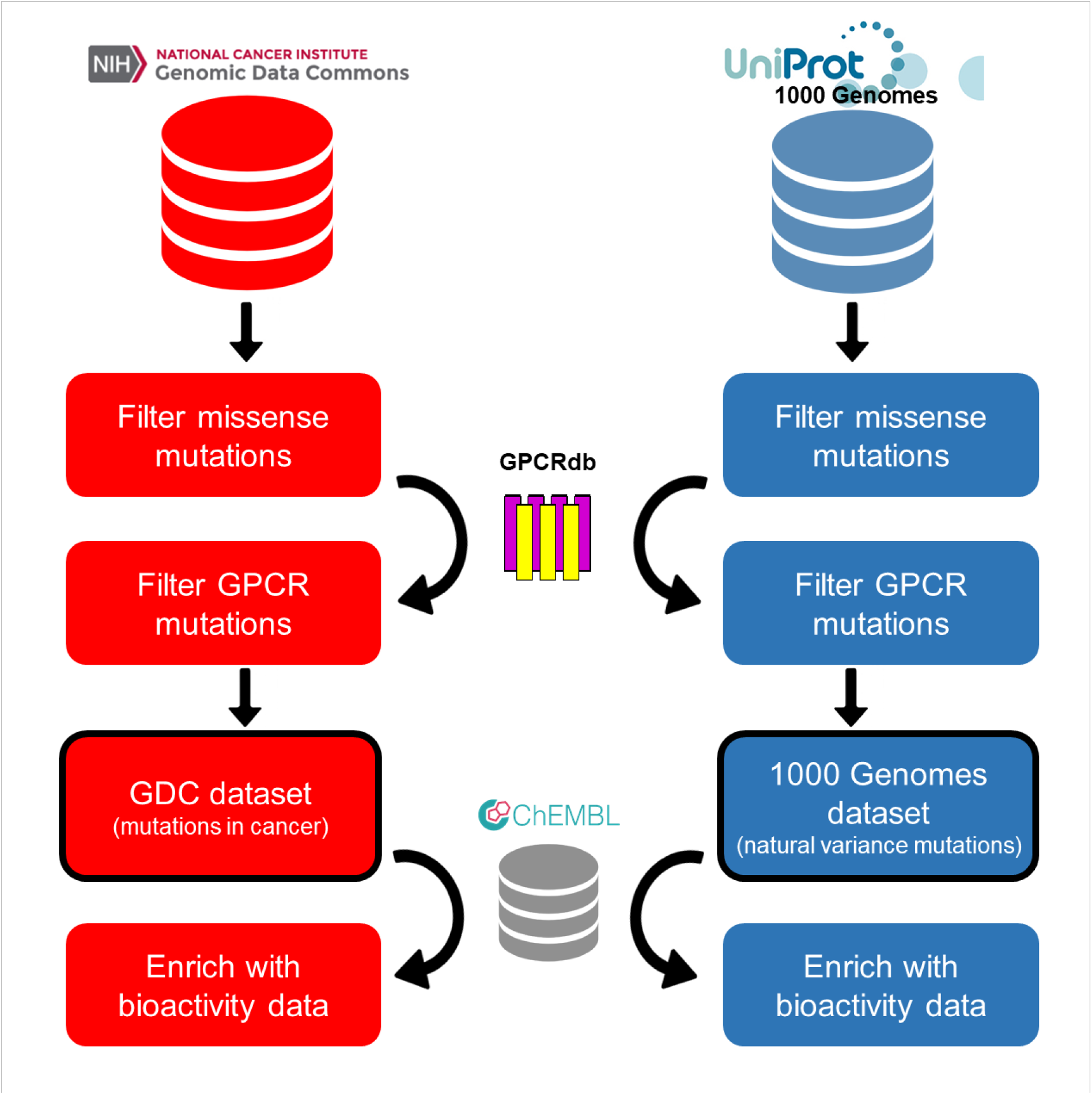
Construction of the GDC and 1000 Genomes datasets. For both the GDC and the Uniprot-1000 Genomes set, missense, and GPCR mutations were filtered, after which identification and annotation data was added from GPCRdb. In a later step, bioactivity data was added from ChEMBL27 for Pareto sorting.

### Bioactivity data

From ChEMBL (release 27) [45,92] ligand-protein interaction data was gathered for all GPCRs in GPCRdb [90]. Data points were retrieved taking into account the following filtering steps: a confidence score of 9, an available pChEMBL value, and the protein belonging to the GPCR-family as defined by the L2 protein class [93]. A pChEMBL value is a standardized value that equals to the negative logarithm of the measured activity for records with dose–response activity types.

### Multiple sequence alignment

The structurally supported alignment provided by GPCRdb was used to study sequence conservation and link sequence positions to the well-established Ballesteros-Weinstein (BW) numbering [12]. A BW analysis can be used to compare positions between GPCRs but is limited to the TM domains. A BW number consists of two parts separated by a decimal sign. The first identifies the TM where this residue is found, and the second number is relative to the most conserved residue in that TM. The most conserved residue is defined to be position 50, with downstream positions receiving a lower number (towards the N-terminus) and upstream positions receiving a higher number (towards the C-terminus). When discrepancies in the BW number were found in the alignment, the most common label was used.

### Two-Entropy Analysis

Two-entropy analysis (TEA) was performed as described previously with slight modifications [19,20]. We started from the TEA algorithm as adjusted by Ye at al. to account for gaps in the multiple sequence alignment as well as for the differences in number of subfamily members [19]. Figure S8 shows the results of our re-implementation in the synthetic dataset provided in [19]. Hereafter, we renamed “Total entropy” as “Rescaled Shannon entropy” and “Average entropy” as “Average entropy across families” for clarification. Firstly, we adapted the previous implementation by using the GPCRdb hierarchy levels to define GPCR subfamilies, resulting in 81 subfamilies for analysis. Secondly, we did not limit the entropy calculation to class A GPCRS but applied it to all GPCRs. However, as opposed to the previous implementation, we included only human GPCR sequences, resulting in 388 sequences for analysis.

### Structural information

The data set was enriched with structural information from GPCRdb [90]. This consisted of data that was annotated to the GPCRs present in the GDC and 1000 Genomes dataset. Included were the family trees to find related proteins, the amino acid sequence of a protein and sequence alignment data, which was used to add BW numbering to the residues. Finally, to connect all the data we found, we used the HUGO Gene Nomenclature Committee (HGNC) for source to source mapping [94].

### Investigated motifs

Several conserved motifs commonly found in GPCRs were further investigated. The following motifs and their surrounding residues, three downstream and three upstream, were investigated (Table 2).

**Table 2:** Investigated motifs, and their residues as noted by their Ballesteros-Weinstein numbering.

| Motif | Residues (Ballesteros-Weinstein number) |
| --- | --- |
| DRY | 3.49, 3.50, 3.51 |
| DRY extended | 3.46 - 3.54 |
| CWxP | 6.47, 6.48, 6.49, 6.50 |
| CWxP extended | 6.44 - 6.53 |
| NPxxY | 7.49, 7.50, 7.51, 7.52, 7.53 |
| NPxxY extended | 7.46 - 7.56 |

*"Extended" in this context refers to the six surrounding residues next to conserved motif.*

### Quantitative structure-activity relationship (QSAR) model training

The performed QSAR models were Random Forest R models trained in Pipeline Pilot using 500 trees and a default seed of “12345” [43,95]. A 50/50 percent training/ hold-out test set was used in duplicate to create and validate these models, with ECFP6 used as molecular descriptors [46].

### Pareto front

Multi-objective ranking was done within the Pareto method as implemented in Pipeline Pilot (version 18.1) [95]. The following properties were used: Mutation rate in TM domains in GDC (maximized), mutation rate in TM domains in the 1000 Genomes set (minimized), average R^2^ of ChEMBL QSAR prediction models (maximized), and in-house availability for experimental assays (maximized). The first and second front were used in further analysis, but all data is provided as supporting information.

### 3D Analysis

*CCR5* crystal structure (PDB code 4MBS) was obtained from the Protein Data Bank [47]. Mutagenesis data was retrieved from the GPCRdb[90] and mapped onto the 3D crystal structure using PyMol [96].

### Hardware

Sequence analysis, data processing, and QSAR modeling were run on a Linux server running CentOS 7. The server had the following components: 2x Intel Xeon Platinum 8160 (2.10), 48 cores, 256 DDR3 RAM, the jobs directory was located on a 1.6 TB PCIe SSD.

### Software

Accelrys Pipeline Pilot 2018 (version 18) was used for all the calculations and analysis [95]. Any calculations performed were done in SI units, using the infrastructure provided in Pipeline Pilot. Data was written towards plain text files and Excel. Graphs were created using Python’s module Matplotlib [97].

## Acknowledgements

The results shown here are in part based upon data generated by the TCGA Research Network: https://www.cancer.gov/tcga. This work was funded by the Dutch Research Council domain Applied and Engineering Sciences (AES) for financial support (STW-Veni #14410). MGG was supported by ONCODE funding. The authors declare that they have no competing interests.

## Data availability statement

The datasets and analysis code supporting the conclusions of this article are available in the 4TU repository DOI: 10.4121/15022410, including the mySQL GDC implementation. The source code used to produce the results in this manuscript was generated in the commercial software package Accelrys Pipeline Pilot 2018 (version 18). The 1000 Genomes dataset was obtained from UNIPROT ‘ftp://ftp.uniprot.org/pub/databases/uniprot/current_release/knowledgebase/variants/’. ChEMBL data was obtained from ‘https://www.ebi.ac.uk/chembl/’, GPCRdb data was obtained from ‘https://gpcrdb.org/’ and HGNC mapping data was obtained from ‘https://www.genenames.org/’.

## References

1. Wild CP, Weiderpass E, Stewart BW. World Cancer Report: Cancer Research for Cancer Prevention. 2020.

2. Bar-Shavit R, Maoz M, Kancharla A, Nag J, Agranovich D, Grisaru-Granovsky S, et al. G Protein-Coupled Receptors in Cancer. Int J Mol Sci. 2016;17: 1320. doi:10.3390/ijms17081320

3. Eckhouse S, Lewison G, Sullivan R. Trends in the global funding and activity of cancer research. Mol Oncol. 2008;2: 20–32. doi:10.1016/j.molonc.2008.03.007

4. Tomczak K, Czerwińska P, Wiznerowicz M. The Cancer Genome Atlas (TCGA): an immeasurable source of knowledge. Contemp Oncol. 2015;19: A68. doi:10.5114/wo.2014.47136

5. Knudson AG. Two genetic hits (more or less) to cancer. Nat Rev Cancer. 2001;1: 157–162. doi:10.1038/35101031

6. O’Hayre M, Vázquez-Prado J, Kufareva I, Stawiski EW, Handel TM, Seshagiri S, et al. The emerging mutational landscape of G proteins and G-protein-coupled receptors in cancer. Nat Rev Cancer. 2013;13: 412–424. doi:10.1038/nrc3521

7. O’Hayre M, Degese MS, Gutkind JS. Novel insights into G protein and G protein-coupled receptor signaling in cancer. Curr Opin Cell Biol. 2014;27: 126–135. doi:10.1016/j.ceb.2014.01.005

8. Hauser AS, Chavali S, Masuho I, Jahn LJ, Martemyanov KA, Gloriam DE, et al. Pharmacogenomics of GPCR Drug Targets. Cell. 2018;172: 41–54.e19. doi:10.1016/j.cell.2017.11.033

9. Peeters MC, van Westen GJP, Li Q, IJzerman AP. Importance of the extracellular loops in G protein-coupled receptors for ligand recognition and receptor activation. Trends Pharmacol Sci. 2011;32: 35–42. doi:10.1016/J.TIPS.2010.10.001

10. Cvicek V, Goddard WA, Abrol R. Structure-Based Sequence Alignment of the Transmembrane Domains of All Human GPCRs: Phylogenetic, Structural and Functional Implications. Ben-Tal N, editor. PLOS Comput Biol. 2016;12: e1004805. doi:10.1371/journal.pcbi.1004805

11. Venkatakrishnan A, Flock T, Prado DE, Oates ME, Gough J, Madan Babu M. Structured and disordered facets of the GPCR fold. Curr Opin Struct Biol. 2014;27: 129–137. doi:10.1016/j.sbi.2014.08.002

12. Ballesteros JA, Weinstein H. [19] Integrated methods for the construction of three-dimensional models and computational probing of structure-function relations in G protein-coupled receptors. 1995. pp. 366–428. doi:10.1016/S1043-9471(05)80049-7

13. Thorpe LM, Yuzugullu H, Zhao JJ. PI3K in cancer: divergent roles of isoforms, modes of activation and therapeutic targeting. Nat Rev Cancer. 2015;15: 7–24. doi:10.1038/nrc3860

14. Liu P, Cheng H, Roberts TM, Zhao JJ. Targeting the phosphoinositide 3-kinase pathway in cancer. Nat Rev Drug Discov. 2009;8: 627–644. doi:10.1038/nrd2926

15. Nairismägi M-L, Tan J, Lim JQ, Nagarajan S, Ng CCY, Rajasegaran V, et al. JAK-STAT and G-protein-coupled receptor signaling pathways are frequently altered in epitheliotropic intestinal T-cell lymphoma. Leukemia. 2016;30: 1311–1319. doi:10.1038/leu.2016.13

16. Pon JR, Marra MA. Driver and Passenger Mutations in Cancer. Annu Rev Pathol Mech Dis. 2015;10: 25–50. doi:10.1146/annurev-pathol-012414-040312

17. Rovati GE, Rie Capra V, Neubig RR. The Highly Conserved DRY Motif of Class A G Protein-Coupled Receptors: Beyond the Ground State. Mol Pharmacol. 2007;71: 959–964. doi:10.1124/mol.106.029470

18. Fritze O, Filipek S, Kuksa V, Palczewski K, Peter Hofmann K, Ernst OP. Role of the conserved NPxxY(x) 5,6 F motif in the rhodopsin ground state and during activation.

19. Ye K, Vriend G, IJzerman AP. Tracing evolutionary pressure. Bioinformatics. 2008;24: 908–915. doi:10.1093/bioinformatics/btn057

20. Ye K, Lameijer EWM, Beukers MW, Ijzerman AP. A two-entropies analysis to identify functional positions in the transmembrane region of class A G protein-coupled receptors. Proteins Struct Funct Genet. 2006;63: 1018–1030. doi:10.1002/prot.20899

21. Erdogmus S, Storch U, Danner L, Becker J, Winter M, Ziegler N, et al. Helix 8 is the essential structural motif of mechanosensitive GPCRs. Nat Commun. 2019;10. doi:10.1038/s41467-019-13722-0

22. Abreu AP, Noel SD, Xu S, Carroll RS, Latronico AC, Kaiser UB. Evidence of the importance of the first intracellular loop of prokineticin receptor 2 in receptor function. Mol Endocrinol. 2012;26: 1417–1427. doi:10.1210/me.2012-1102

23. Connolly A, Holleran BJ, Simard É, Baillargeon JP, Lavigne P, Leduc R. Interplay between intracellular loop 1 and helix VIII of the angiotensin II type 2 receptor controls its activation. Biochem Pharmacol. 2019;168: 330–338. doi:10.1016/j.bcp.2019.07.018

24. Wang ZQ, Tao YX. Functional studies on twenty novel naturally occurring melanocortin-4 receptor mutations. Biochim Biophys Acta - Mol Basis Dis. 2011;1812: 1190–1199. doi:10.1016/j.bbadis.2011.06.008

25. Johnson MS, Robertson DN, Holland PJ, Lutz EM, Mitchell R. Role of the conserved NPxxY motif of the 5-HT2A receptor in determining selective interaction with isoforms of ADP-Ribosylation Factor (ARF). Cell Signal. 2006;18: 1793–1800. doi:10.1016/J.CELLSIG.2006.02.002

26. Kim J-M, Altenbach C, Thurmond RL, Khorana HG, Hubbell WL, Ernst OP. Structure and function in rhodopsin: Rhodopsin mutants with a neutral amino acid at E134 have a partially activated conformation in the dark state. Proc Natl Acad Sci. 1997;94: 14273–14278. doi:10.1073/pnas.94.26.14273

27. Olivella M, Caltabiano G, Cordomí A. The role of Cysteine 6.47 in class A GPCRs. BMC Struct Biol. 2013;13: 3. doi:10.1186/1472-6807-13-3

28. Nomiyama H, Yoshie O. Functional roles of evolutionary conserved motifs and residues in vertebrate chemokine receptors. J Leukoc Biol. 2015;97: 39–47. doi:10.1189/jlb.2RU0614-290R

29. Chung DA, Wade SM, Fowler CB, Woods DD, Abada PB, Mosberg HI, et al. Mutagenesis and peptide analysis of the DRY motif in the α2A adrenergic receptor: evidence for alternate mechanisms in G protein-coupled receptors. Biochem Biophys Res Commun. 2002;293: 1233–1241. doi:10.1016/S0006-291X(02)00357-1

30. Rasmussen SGF, Jensen AD, Liapakis G, Ghanouni P, Javitch JA, Gether U, et al. Mutation of a Highly Conserved Aspartic Acid in the β_2_ Adrenergic Receptor: Constitutive Activation, Structural Instability, and Conformational Rearrangement of Transmembrane Segment 6. Mol Pharmacol. 1999;56: 175–184. doi:10.1124/mol.56.1.175

31. Kim K-M, Caron MG. Complementary roles of the DRY motif and C-terminus tail of GPCRS for G protein coupling and β-arrestin interaction. Biochem Biophys Res Commun. 2008;366: 42–47. doi:10.1016/J.BBRC.2007.11.055

32. Kolakowski LF. GCRDb: A G-protein-coupled receptor database. Receptors and Channels. 1994. pp. 1–7.

33. Shah G V, Rayford W, Noble MJ, Austenfeld M, Weigel J, Vamos S, et al. Calcitonin stimulates growth of human prostate cancer cells through receptor-mediated increase in cyclic adenosine 3’,5’-monophosphates and cytoplasmic Ca2+ transients. Endocrinology. 1994;134: 596–602. doi:10.1210/endo.134.2.8299557

34. Gillespie MT, Thomas RJ, Pu Z-Y, Zhou H, Martin TJ, Findlay DM. Calcitonin receptors, bone sialoprotein and osteopontin are expressed in primary breast cancers. Int J Cancer. 1997;73: 812–815. doi:10.1002/(SICI)1097-0215(19971210)73:6<812::AID-IJC7>3.0.CO;2-5

35. Benes L, Kappus C, McGregor GP, Bertalanffy H, Mennel HD, Hagner S. The immunohistochemical expression of calcitonin receptor-like receptor (CRLR) in human gliomas. J Clin Pathol. 2004;57: 172–176. doi:10.1136/jcp.2003.12997

36. Nikitenko LL, Leek R, Henderson S, Pillay N, Turley H, Generali D, et al. The G-protein-coupled receptor CLR is upregulated in an autocrine loop with adrenomedullin in clear cell renal cell carcinoma and associated with poor prognosis. Clin Cancer Res. 2013;19: 5740–5748. doi:10.1158/1078-0432.CCR-13-1712

37. Acconcia F. The Network of Angiotensin Receptors in Breast Cancer. Cells. 2020;9: 1–14. doi:10.3390/cells9061336

38. Hill SM, Frasch T, Xiang S, Yuan L, Duplessis T, Mao L. Molecular mechanisms of melatonin anticancer effects. Integr Cancer Ther. 2009;8: 337–346. doi:10.1177/1534735409353332

39. Sarrouilhe D, Mesnil M. Serotonin and human cancer: A critical view. Biochimie. 2019;161: 46–50. doi:10.1016/j.biochi.2018.06.016

40. Teh J, Chen S. mGlu Receptors and Cancerous Growth. Wiley Interdiscip Rev Membr Transp Signal. 2011;1: 211–220. doi:10.1002/wmts.21.mGlu

41. Fröhlich E, Wahl R. The forgotten effects of thyrotropin-releasing hormone: Metabolic functions and medical applications. Front Neuroendocrinol. 2019;52: 29–43. doi:10.1016/j.yfrne.2018.06.006

42. Kamath J, Feinn R, Winokur A. Thyrotropin-releasing hormone as a treatment for cancer-related fatigue: A randomized controlled study. Support Care Cancer. 2012;20: 1745–1753. doi:10.1007/s00520-011-1268-8

43. Svetnik V, Liaw A, Tong C, Culberson JC, Sheridan RP, Feuston BP. Random forest: a classification and regression tool for compound classification and QSAR modeling. J Chem Inf Comput Sci. 2003;43: 1947–58. doi:10.1021/ci034160g

44. Gaulton A, Hersey A, -l Nowotka M, Patrícia Bento A, Chambers J, Mendez D, et al. The ChEMBL database in 2017. Nucleic Acids Res. 2016;45: 945–954. doi:10.1093/nar/gkw1074

45. ChEMBL27 Database Release. doi:10.6019/CHEMBL.database.27

46. Wang X, Jespers W, Bongers BJ, Habben Jansen MCC, Stangenberger CM, Dilweg MA, et al. Characterization of cancer-related somatic mutations in the adenosine A2B receptor. Eur J Pharmacol. 2020;880: 173126. doi:10.1016/j.ejphar.2020.173126

47. Tan Q, Zhu Y, Li JJ, Chen Z, Han GW, Kufareva I, et al. Structure of the CCR5 chemokine receptor-HIV entry inhibitor maraviroc complex. Science. 2013;341: 1387–90. doi:10.1126/science.1241475

48. Kleist AB, Getschman AE, Ziarek JJ, Nevins AM, Gauthier P-A, Chevigne A, et al. New paradigms in chemokine receptor signal transduction: moving beyond the two-site model. Biochem Pharmacol. 2016;114: 53–68. doi:10.1016/j.bcp.2016.04.007.New

49. Tan Q, Zhu Y, Li J, Chen Z, Han GW, Kufareva I, et al. Structure of the CCR5 chemokine receptor-HIV entry inhibitor maraviroc complex. Science (80- ). 2013;341: 1387–1390. doi:10.1126/science.1241475

50. Semack A, Sandhu M, Malik RU, Vaidehi N, Sivaramakrishnan S. Structural elements in the Gαs and Gβq C termini that mediate selective G Protein-coupled Receptor (GPCR) signaling. J Biol Chem. 2016;291: 17929–17940. doi:10.1074/jbc.M116.735720

51. Lindner D, Walther C, Tennemann A, Beck-Sickinger AG. Functional role of the extracellular N-terminal domain of neuropeptide Y subfamily receptors in membrane integration and agonist-stimulated internalization. Cell Signal. 2009;21: 61–68. doi:10.1016/j.cellsig.2008.09.007

52. Zhang Z, Wu J, Yu J, Xiao J. A brief review on the evolution of GPCR: conservation and diversification. Open J Genet. 2012;2: 11–17.

53. Tao YX, Segaloff DL. Functional analyses of melanocortin-4 receptor mutations identified from patients with binge eating disorder and nonobese or obese subjects. J Clin Endocrinol Metab. 2005;90: 5632–5638. doi:10.1210/jc.2005-0519

54. Fredriksson R, Lagerström MC, Lundin LG, Schiöth HB. The G-protein-coupled receptors in the human genome form five main families. Phylogenetic analysis, paralogon groups, and fingerprints. Mol Pharmacol. 2003;63: 1256–1272. doi:10.1124/mol.63.6.1256

55. Stoy H, Gurevich V V. How genetic errors in GPCRs affect their function: Possible therapeutic strategies. Genes and Diseases. Chongqing yi ke da xue, di 2 lin chuang xue yuan Bing du xing gan yan yan jiu suo; 2015. pp. 108–132. doi:10.1016/j.gendis.2015.02.005

56. Lagane B, Ballet S, Planchenault T, Balabanian K, Le Poul E, Blanpain C, et al. Mutation of the DRY motif reveals different structural requirements for the CC chemokine receptor 5-mediated signaling and receptor endocytosis. Mol Pharmacol. 2005;67: 1966–76. doi:10.1124/mol.104.009779

57. Zhou Q, Yang D, Wu M, Guo Y, Guo W, Zhong L, et al. Common activation mechanism of class a GPCRs. Elife. 2019;8: 1–31. doi:10.7554/eLife.50279

58. Chen A, Gao ZG, Barak D, Liang BT, Jacobson KA. Constitutive activation of A3 adenosine receptors by site-directed mutagenesis. Biochem Biophys Res Commun. 2001;284: 596–601. doi:10.1006/bbrc.2001.5027

59. Tsunekawa K, Onigata K, Morimura T, Kasahara T, Nishiyama S, Kamoda T, et al. Identification and functional analysis of novel inactivating thyrotropin receptor mutations in patients with thyrotropin resistance. Thyroid. 2006;16: 471–479. doi:10.1089/thy.2006.16.471

60. Jensen PC, Nygaard R, Thiele S, Elder A, Zhu G, Kolbeck R, et al. Molecular interaction of a potent nonpeptide agonist with the chemokine receptor CCR8. Mol Pharmacol. 2008;74: 539. doi:10.1124/mol.107.035543

61. Schlecker E, Stojanovic A, Eisen C, Quack C, Falk CS, Umansky V, et al. Tumor-infiltrating monocytic myeloid-derived suppressor cells mediate CCR5-dependent recruitment of regulatory T cells favoring tumor growth. J Immunol. 2012;189: 5602–11. doi:10.4049/jimmunol.1201018

62. Schmall A, Al-Tamari HM, Herold S, Kampschulte M, Weigert A, Wietelmann A, et al. Macrophage and cancer cell cross-talk via CCR2 and CX3CR1 is a fundamental mechanism driving lung cancer. Am J Respir Crit Care Med. 2015;191: 437–447. doi:10.1164/rccm.201406-1137OC

63. Hao Q, Vadgama J V., Wang P. CCL2/CCR2 signaling in cancer pathogenesis. Cell Commun Signal. 2020;18: 1–13. doi:10.1186/s12964-020-00589-8

64. Houben AJS, Moolenaar WH. Autotaxin and LPA receptor signaling in cancer. Cancer Metastasis Rev. 2011;30: 557–565. doi:10.1007/s10555-011-9319-7

65. Yang L, Lindholm K, Konishi Y, Li R, Shen Y. Target depletion of distinct tumor necrosis factor receptor subtypes reveals hippocampal neuron death and survival through different signal transduction pathways. J Neurosci. 2002;22: 3025–32. doi:20026317

66. Prickett TD, Samuels Y. Molecular Pathways: Dysregulated Glutamatergic Signaling Pathways in Cancer. Clin Cancer Res. 2012;18: 4240–4246. doi:10.1158/1078-0432.CCR-11-1217

67. Zhang P, Kang B, Xie G, Li S, Gu Y, Shen Y, et al. Genomic sequencing and editing revealed the GRM8 signaling pathway as potential therapeutic targets of squamous cell lung cancer. Cancer Lett. 2019;442: 53–67. doi:10.1016/j.canlet.2018.10.035

68. Gwynne WD, Shakeel MS, Girgis-Gabardo A, Kim KH, Ford E, Dvorkin-Gheva A, et al. Antagonists of the serotonin receptor 5A target human breast tumor initiating cells. BMC Cancer. 2020;20: 1–17. doi:10.1186/s12885-020-07193-6

69. Allard B, Cousineau I, Allard D, Buisseret L, Pommey S, Chrobak P, et al. Adenosine A2a receptor promotes lymphangiogenesis and lymph node metastasis. Oncoimmunology. 2019;8: 1–16. doi:10.1080/2162402X.2019.1601481

70. Masjedi A, Ahmadi A, Ghani S, Malakotikhah F, Nabi Afjadi M, Irandoust M, et al. Silencing adenosine A2a receptor enhances dendritic cell-based cancer immunotherapy. Nanomedicine Nanotechnology, Biol Med. 2020;29: 102240. doi:10.1016/j.nano.2020.102240

71. Liu H, Kuang X, Zhang Y, Ye Y, Li J, Liang L, et al. ADORA1 Inhibition Promotes Tumor Immune Evasion by Regulating the ATF3-PD-L1 Axis. Cancer Cell. 2020;37: 324–339.e8. doi:10.1016/j.ccell.2020.02.006

72. Ni S, Wei Q, Yang L. Adora1 promotes hepatocellular carcinoma progression via pi3k/akt pathway. Onco Targets Ther. 2020;13: 12409–12419. doi:10.2147/OTT.S272621

73. Hwang SM, Kim HJ, Kim SM, Jung Y, Park SW, Chung IY. Lysophosphatidylserine receptor P2Y10: A G protein-coupled receptor that mediates eosinophil degranulation. Clin Exp Allergy. 2018;48: 990–999. doi:10.1111/cea.13162

74. Blanpain C, Doranz BJ, Bondue A, Govaerts C, De Leener A, Vassart G, et al. The Core Domain of Chemokines Binds CCR5 Extracellular Domains while Their Amino Terminus Interacts with the Transmembrane Helix Bundle. J Biol Chem. 2003;278: 5179–5187. doi:10.1074/jbc.M205684200

75. Dragic T, Trkola A, Lin SW, Nagashima KA, Kajumo F, Zhao L, et al. Amino-Terminal Substitutions in the CCR5 Coreceptor Impair gp120 Binding and Human Immunodeficiency Virus Type 1 Entry. J Virol. 1998;72: 279–285. doi:10.1128/jvi.72.1.279-285.1998

76. Garcia-Perez J, Rueda P, Alcami J, Rognan D, Arenzana-Seisdedos F, Lagane B, et al. Allosteric model of maraviroc binding to CC chemokine receptor 5 (CCR5). J Biol Chem. 2011;286: 33409–21. doi:10.1074/jbc.M111.279596

77. Jespers W, Schiedel AC, Heitman LH, Cooke RM, Kleene L, van Westen GJP, et al. Structural Mapping of Adenosine Receptor Mutations: Ligand Binding and Signaling Mechanisms. Trends Pharmacol Sci. 2017. doi:10.1016/j.tips.2017.11.001

78. Farzan M, Choe H, Martin KA, Sun Y, Sidelko M, Mackay CR, et al. HIV-1 entry and macrophage inflammatory protein-1beta-mediated signaling are independent functions of the chemokine receptor CCR5. J Biol Chem. 1997;272: 6854–7. doi:10.1074/jbc.272.11.6854

79. Kondru R, Zhang J, Ji C, Mirzadegan T, Rotstein D, Sankuratri S, et al. Molecular interactions of CCR5 with major classes of small-molecule anti-HIV CCR5 antagonists. Mol Pharmacol. 2008;73: 789–800. doi:10.1124/mol.107.042101

80. Swinney DC, Beavis P, Chuang KT, Zheng Y, Lee I, Gee P, et al. A study of the molecular mechanism of binding kinetics and long residence times of human CCR5 receptor small molecule allosteric ligands. Br J Pharmacol. 2014;171: 3364–3375. doi:10.1111/bph.12683

81. Wu V, Yeerna H, Nohata N, Chiou J, Harismendy O, Raimondi F, et al. Illuminating the Onco-GPCRome: Novel G protein-coupled receptor-driven oncocrine networks and targets for cancer immunotherapy. J Biol Chem. 2019;294: 11062–11086. doi:10.1074/jbc.REV119.005601

82. Sriram K, Moyung K, Corriden R, Carter H, Insel PA. GPCRs show widespread differential mRNA expression and frequent mutation and copy number variation in solid tumors. PLoS Biology. 2019. doi:10.1371/journal.pbio.3000434

83. Arakaki AKS, Pan WA, Trejo JA. GPCRs in cancer: Protease-activated receptors, endocytic adaptors and signaling. Int J Mol Sci. 2018;19. doi:10.3390/ijms19071886

84. Jensen MA, Ferretti V, Grossman RL, Staudt LM. The NCI Genomic Data Commons as an engine for precision medicine. Blood. 2017;130: 453–459. doi:10.1182/blood-2017-03-735654

85. Broad Institute of MIT and Harvard. Firehose 2015_11_01 run. 2015. doi:10.7908/C1571BB1

86. Axmark D, Widenius M. MySQL 5.7 reference manual. Redwood Shores, CA: Oracle. 2015. Available: http://dev.mysql.com/doc/refman/5.7/en/index.html

87. Auton A, Abecasis GR, Altshuler DM, Durbin RM, Abecasis GR, Bentley DR, et al. A global reference for human genetic variation. Nature. 2015;526: 68–74. doi:10.1038/nature15393

88. The UniProt Consortium. UniProt: a worldwide hub of protein knowledge. Nucleic Acids Res. 2019;47. doi:10.1093/nar/gky1049

89. Tate JG, Bamford S, Jubb HC, Sondka Z, Beare DM, Bindal N, et al. COSMIC: the Catalogue Of Somatic Mutations In Cancer. Nucleic Acids Res. 2019;47: D941–D947. doi:10.1093/nar/gky1015

90. Isberg V, Mordalski S, Munk C, Rataj K, Harpsoe K, Hauser AS, et al. GPCRdb: An information system for G protein-coupled receptors. Nucleic Acids Res. 2016;44: D356–D364. doi:10.1093/nar/gkv1178

91. Finan C, Gaulton A, Kruger FA, Lumbers RT, Shah T, Engmann J, et al. The druggable genome and support for target identification and validation in drug development. Sci Transl Med. 2017;9. doi:10.1126/scitranslmed.aag1166

92. Gaulton A, Hersey A, Nowotka ML, Patricia Bento A, Chambers J, Mendez D, et al. The ChEMBL database in 2017. Nucleic Acids Res. 2017;45: D945–D954. doi:10.1093/nar/gkw1074

93. Papadatos G, Gaulton A, Hersey A, Overington JP. Activity, assay and target data curation and quality in the ChEMBL database. J Comput Aided Mol Des. 2015;29: 885–96. doi:10.1007/s10822-015-9860-5

94. Bruford EA, Lush MJ, Wright MW, Sneddon TP, Povey S, Birney E. The HGNC Database in 2008: a resource for the human genome. Nucleic Acids Res. 2007;36: D445–D448. doi:10.1093/nar/gkm881

95. BIOVIA Pipeline Pilot | Scientific Workflow Authoring Application for Data Analysis.

96. The PyMOL Molecular Graphics System, Version 1.4 Schrödinger, LLC.

97. Hunter JD. Matplotlib: A 2D Graphics Environment. Comput Sci Eng. 2007;9: 90–95. doi:10.1109/MCSE.2007.55

