## Supplementary Figures 1-8 for "Pan-cancer in silico analysis of somatic mutations in G-protein coupled receptors: The effect of evolutionary conservation and natural variance"

### Supplementary Information

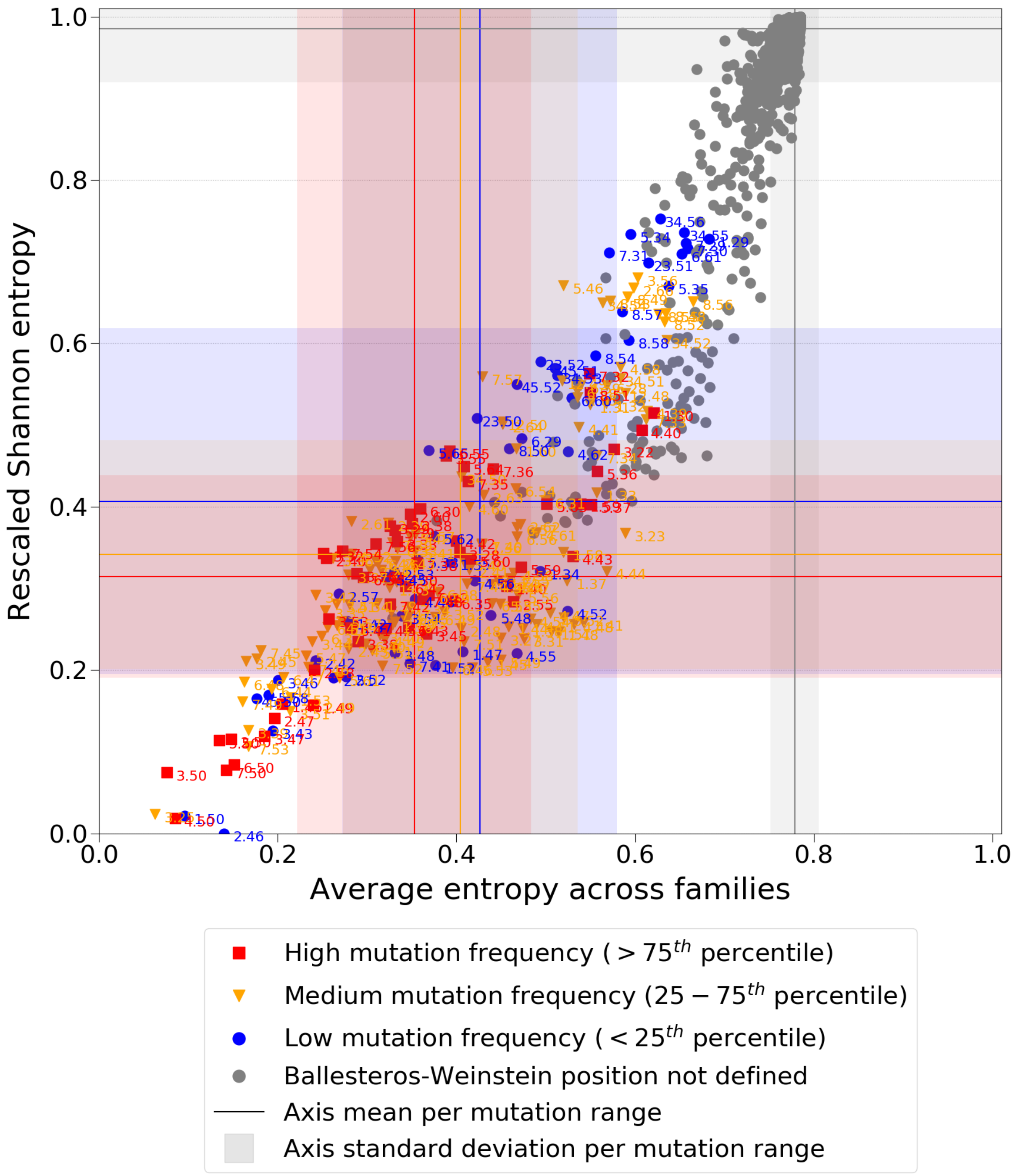


***Fig S1: Residue Shannon entropy, with residue and GDC labels.*** *A two-entropy analysis plot for all GPCRs with aligned positions and labelled residues. The average entropy across families, i.e. conserved within a family is on the x-axis, and the Shannon entropy overall on the y-axis. Residues are colored by the frequency of mutations found in the GDC dataset, with blue being low (< 25th percentile), orange medium (25-75th percentiles) and red high (> 75th percentile). Residues with no defined Ballesteros-Weinstein labels are colored grey. Blue, orange, red, and grey lines represent the mean entropy values for each axis per mutation range (high, medium, low, and non-defined Ballesteros-Weinstein, respectively). Blue, orange, red, and grey shadows represent the standard deviation to the mean entropy values for each axis per mutation range (high, medium, low, and non-defined Ballesteros-Weinstein, respectively).*

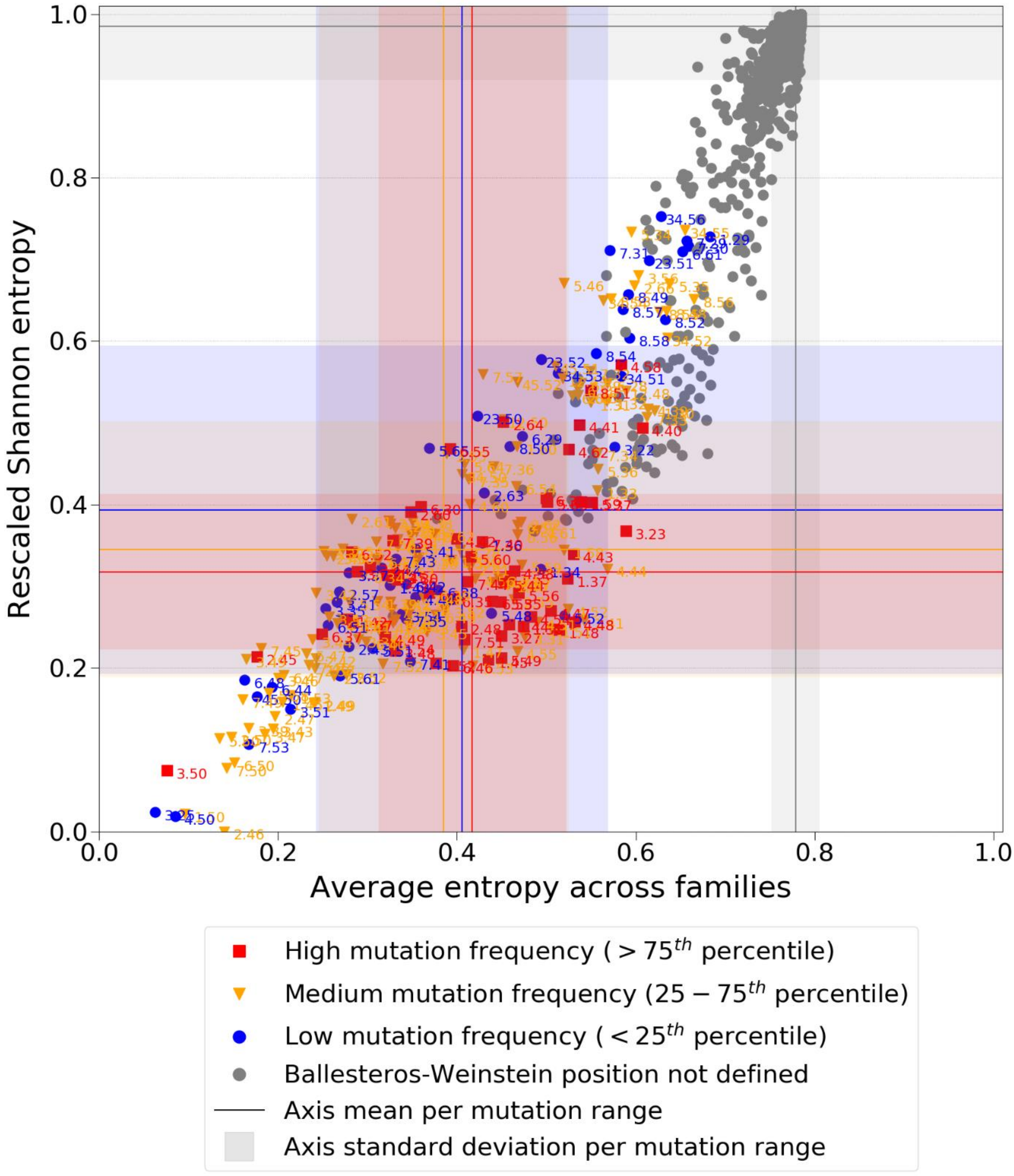


***Fig S2: Residue Shannon entropy, with residue and 1000 Genomes labels.*** *A two-entropy analysis plot for all GPCRs with aligned positions and labelled residues. The average entropy across families, i.e. conserved within a family is on the x-axis, and the Shannon entropy overall on the y-axis. Residues are colored by the frequency of mutations found in the 1000 Genomes dataset, with blue being low (< 25$^{th}$ percentile), orange medium (25-75$^{th}$ percentiles) and red high (> 75$^{th}$ percentile). Residues with no defined Ballesteros-Weinstein labels are colored grey. Blue, orange, red, and grey lines represent the mean entropy values for each axis per mutation range (high, medium, low, and non-defined Ballesteros-Weinstein, respectively). Blue, orange, red, and grey shadows represent the standard deviation to the mean entropy values for each axis per mutation range (high, medium, low, and non-defined Ballesteros-Weinstein, respectively).*

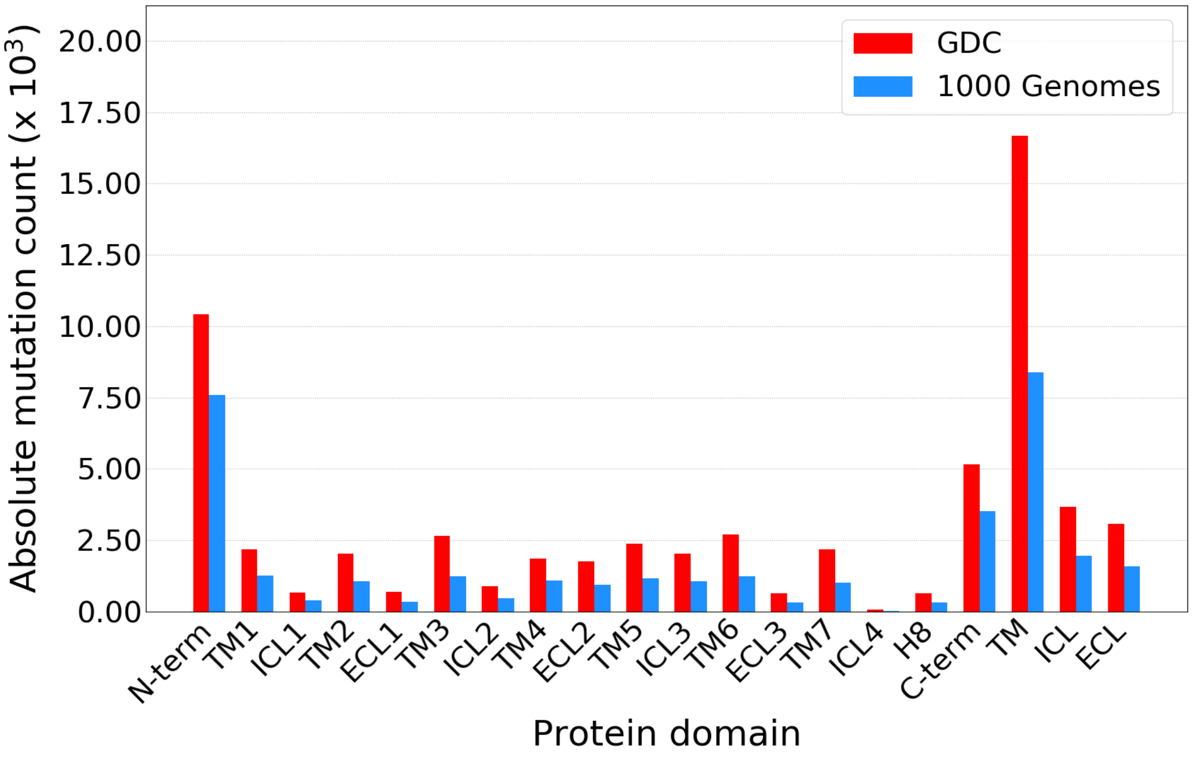


***Fig S3: Absolute mutation count per GPCR protein domain.*** *Absolute mutation count found in the GDC and 1000 Genomes data, split on the domain in GPCRs where they were found in. "TM", "ICL" and "ECL" represent the aggregated fractions for those domains.*

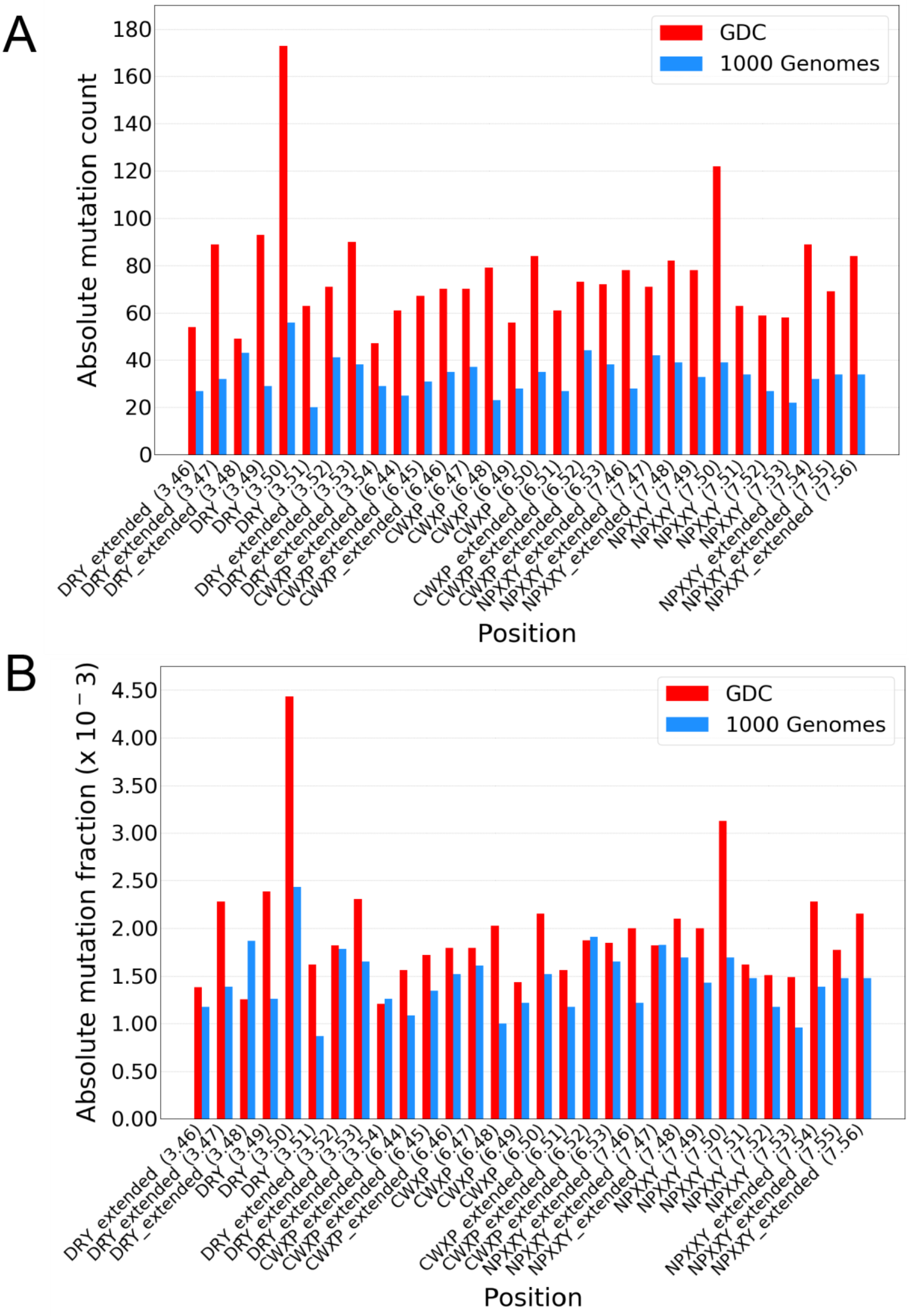


***Fig S4: Mutations per extended conserved motif residue.*** *(A) Absolute mutation count of the GDC and 1000 Genomes set for the extended motifs of "DRY" (TM3), "CWxP" (TM6) and "NPxxY" (TM7). (B) Absolute mutation fraction of the GDC and 1000 Genomes set for the extended motifs of "DRY" (TM3), "CWxP" (TM6) and "NPxxY" (TM7).*

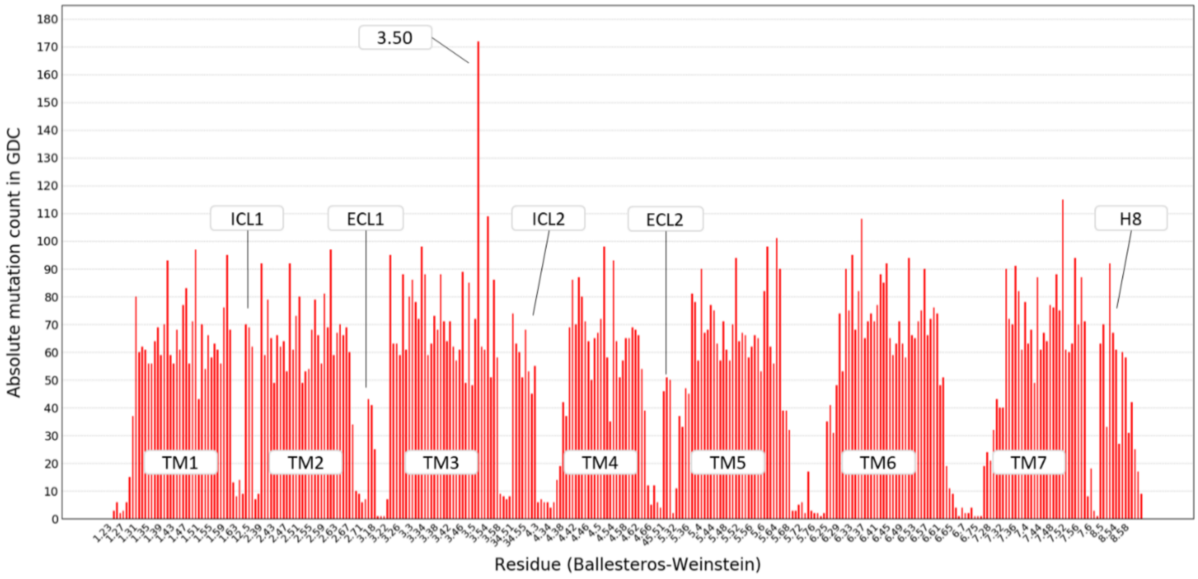


***Fig S5: GPCR cancer mutations on Ballesteros-Weinstein positions.*** *GPCR cancer mutations plotted for the Ballesteros-Weinstein positions found in the GDC data. Positions are ordered from lowest to highest and X-axis labels are displayed every three residues for visualization purposes.*

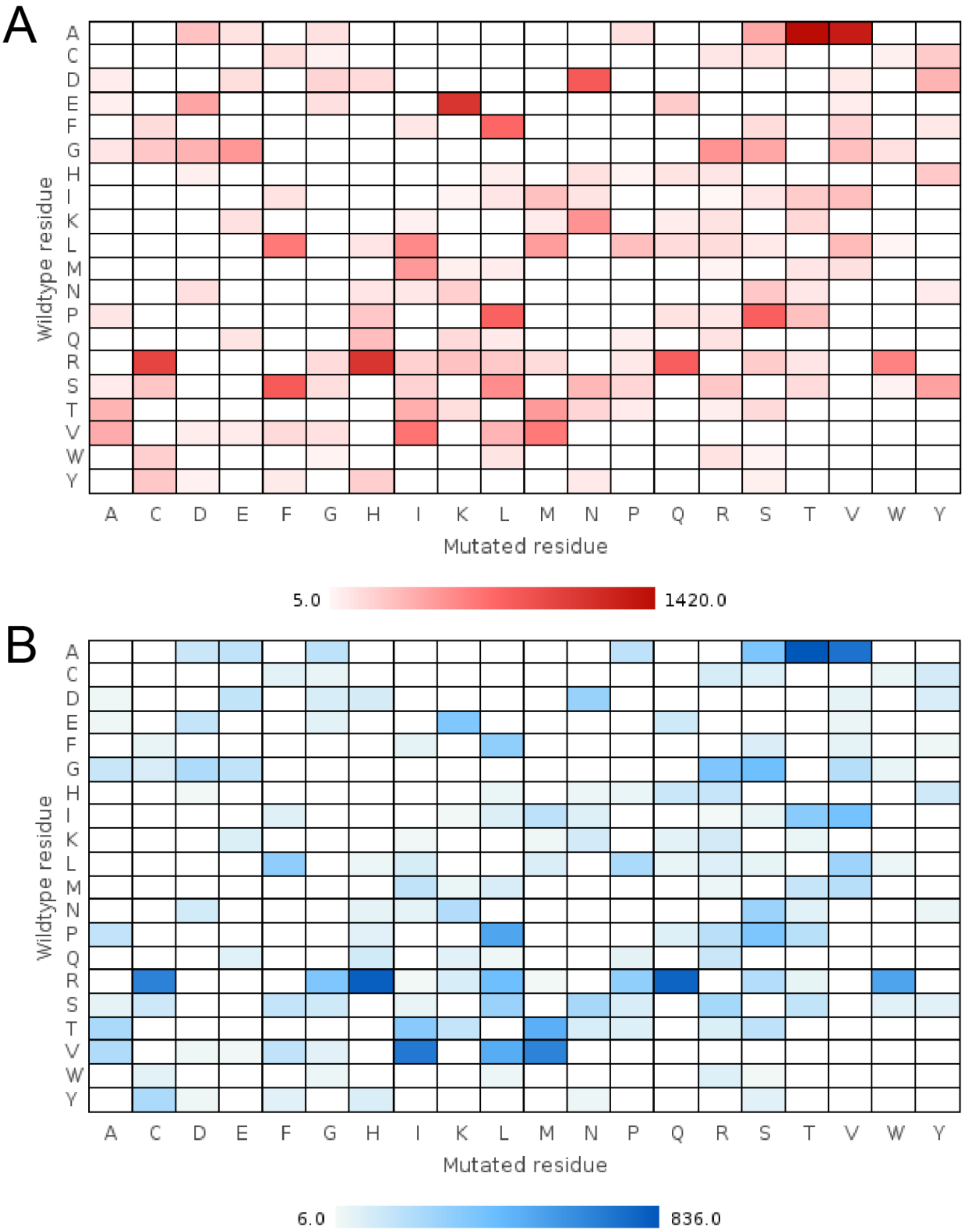


***Fig S6: Heat-map cancer substitutions.*** *(A) Heat-map showing the frequency of substitutions found in the GDC dataset. A darker shade of red means a higher frequency. (B) Heat-map showing the frequency of substitutions found in the 1000 Genomes dataset. A darker shade of blue means a higher frequency.*

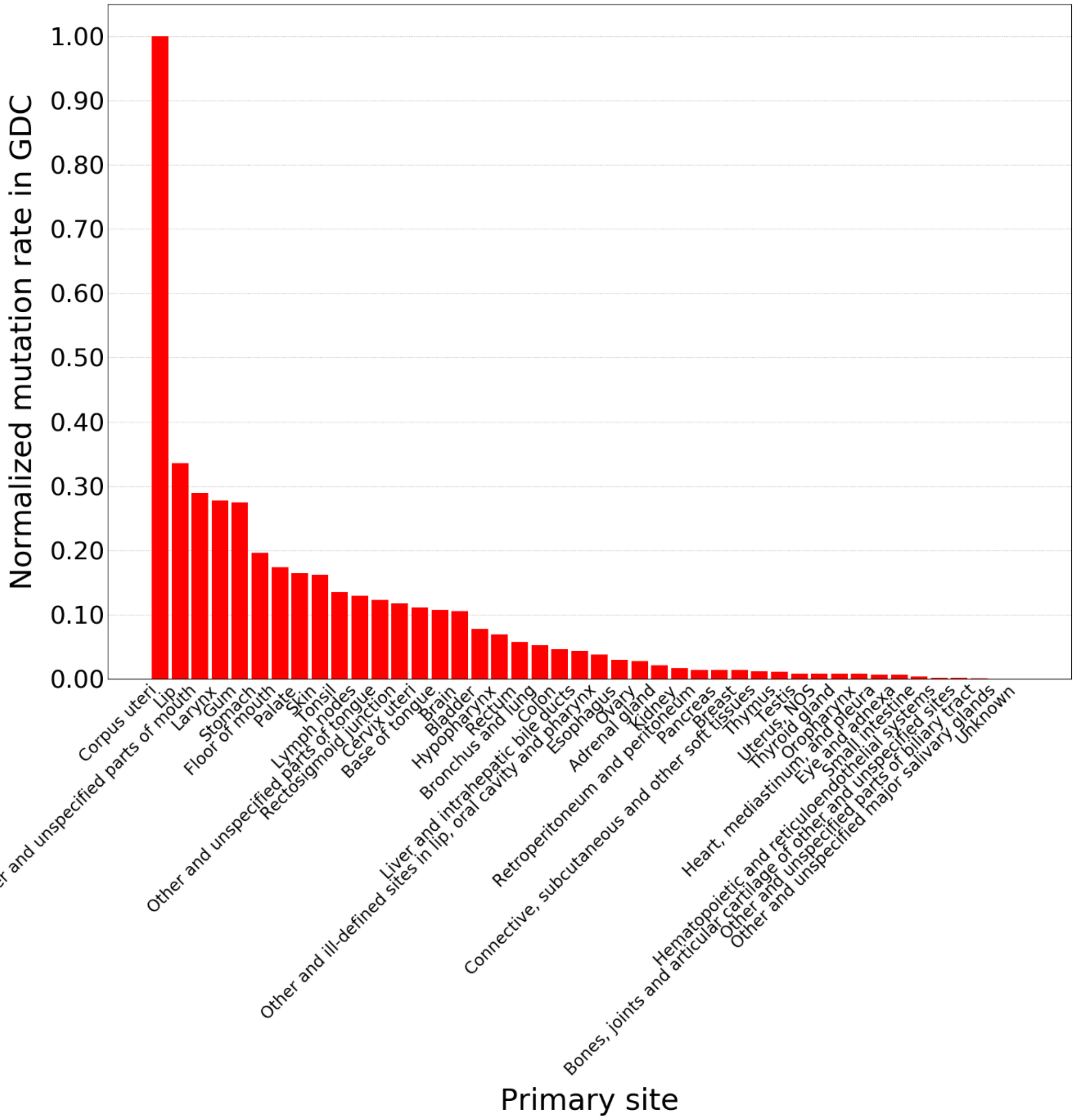


***Fig S7: GPCR mutation rates by cancer type.*** *Normalized GPCR mutation rate per primary site (i.e. cancer type). Mutation rate per primary site is normalized by the number of patients in GDC with that cancer type.*

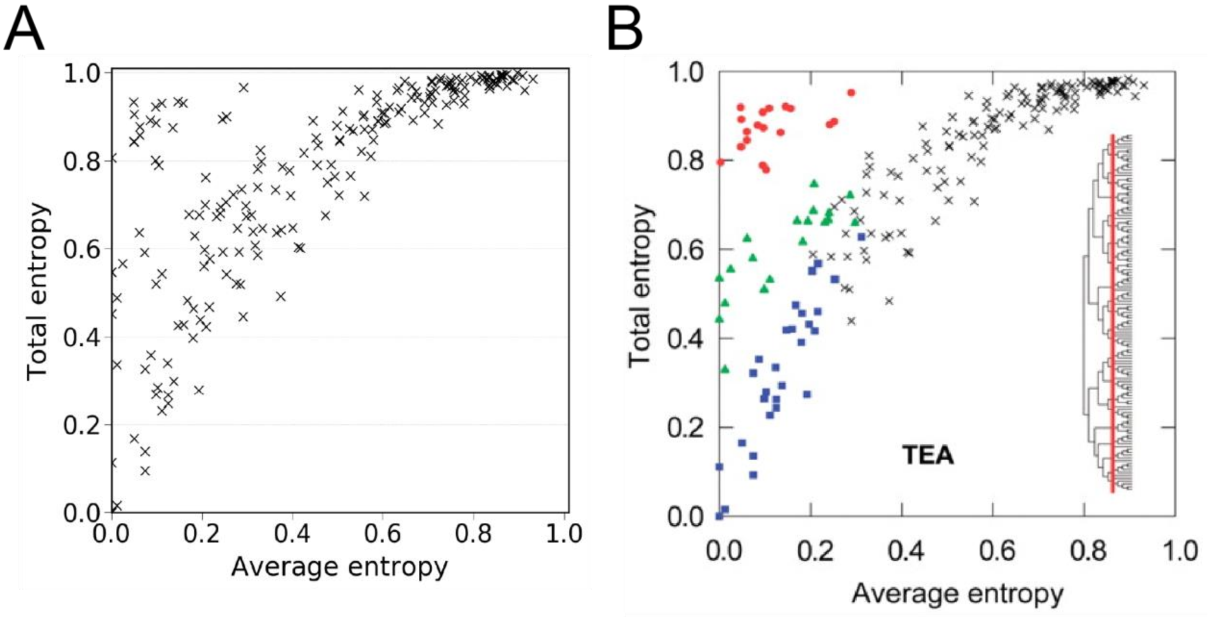


***Fig S8: Two-entropy analysis re-implementation.*** *(A) Re-implementation of two-entropy analysis in a synthetic dataset as defined by Ye et al. in* [19]. *(B) Original analysis, figure adapted from Ye et al. in [19].*
